# Necroptosis is a targetable driver of severe coronavirus pathogenesis

**DOI:** 10.64898/2026.11.09.765133

**Authors:** Yuxin Hou, Jialu Shi, Minmin Zhou, Baokun Sui, Honglei Zhang, Ye-Fan Hu, Haibo Li, Yixin He, Zhen Sun, Zhe Chen, Carly DeAntoneo, Huan Liu, Jianwei Li, Avishekh Gautam, Cuiting Luo, Lei Wen, Chaemin Yoon, Yanxia Chen, Yixin Wang, Tianrenzheng Zhu, Yuanchen Liu, Bingjie Hu, Terrence Tsz-Tai Yuen, Xi Zhang, Jianpiao Cai, Huiping Shuai, Cun Li, Xiaoyu Zhao, Mengyao Liu, Xiancai Ma, Shuofeng Yuan, Jie Zhou, Bao-Zhong Zhang, Kwok Fai Lucius Lee, Zai Wang, Kin-Hang Kok, Jian-Dong Huang, Dong-Yan Jin, Jasper Fuk-Woo Chan, Bin Cao, Francis Ka-Ming Chan, Siddharth Balachandran, Kwok-Yung Yuen, Hin Chu

## Abstract

Human-pathogenic coronaviruses differ dramatically in pathogenicity but the underlying mechanisms are incompletely understood. Here, we demonstrate that highly pathogenic coronaviruses trigger necroptosis in the infected cells and lung tissues, whereas necroptosis is not detected in cells infected by seasonal coronavirus. Mechanistically, highly pathogenic coronaviruses produce Z-RNA, activate ZBP1 and initiates necroptotic cell death signaling, triggering necroinflammation and lung injury. In contrast, seasonal coronaviruses produce little to no Z-RNA, and do not activate necroptosis. Importantly, small-molecule inhibition of necroptosis attenuates inflammatory cytokine expression and markedly ameliorates lung damage in mice infected with all three highly pathogenic coronaviruses, independent of virus clearance. Overall, our study identifies necroptosis as a dominant mediator of coronavirus pathogenesis, and positions necroptosis blockade as a new therapeutic target against severe coronavirus diseases.

**One Sentence Summary:** Necroptosis is a driver of severe coronavirus pathogenesis that is targetable through small-molecule inhibition

## INTRODUCTION

Seven coronaviruses including human coronavirus (HCoV)-229E, HCoV-OC43, severe acute respiratory syndrome (SARS)-CoV-1, HCoV-NL63, HCoV-HKU1, Middle East respiratory syndrome (MERS)-CoV, and SARS-CoV-2, have caused pandemics or epidemics in humans(*1, 2*). More recently, human porcine deltacoronavirus (Hu-PDCoV), a close relative of porcine deltacoronavirus HKU15(*3*), canine coronavirus human pneumonia 2018 (CCoV-HuPn-2018), and rodent alphacoronavirus (HCoV-KUMC22-3) have been identified in humans, highlighting the persistent risk of zoonotic coronavirus spillovers(*4–6*). While infection by highly pathogenic coronaviruses (SARS-CoV-1, MERS-CoV, and SARS-CoV-2) can result in cytokine storm, extensive lung injury, acute respiratory distress syndrome (ARDS) and death, infection by seasonal coronaviruses predominantly result in self-limiting upper respiratory tract infections, with symptoms akin to common colds(*7*). The core mechanisms that govern the virulence of human-pathogenic coronaviruses remain incompletely understood.

A common feature of coronavirus infection is their propensity to induce cell damage in the tissues of infected patients(*8–11*), suggesting the involvement of programmed cell death pathways in coronavirus pathogenesis. In keeping with these observations, we and others recently demonstrated that apoptosis triggered during coronavirus infection promotes virus replication and pathogenesis(*12–14*). In addition to apoptosis, necroptosis is a regulated cell death mechanism(*15–19*) that plays key roles in the biology of both DNA(*20–24*) and RNA(*17, 25–28*) viruses. Necroptosis activation is documented in autopsied lung tissues from patients who succumbed to COVID-19(*29, 30*) but its role in driving pathogenesis during SARS-CoV-2 infection remains inconclusive(*31–35*). Furthermore, it is unclear whether necroptosis is triggered by other coronaviruses, particularly highly virulent coronaviruses, such as MERS-CoV and SARS-CoV-1. Lastly, the mechanism(s) by which coronaviruses trigger necroptosis, and the importance of necroptotic cell death to either antiviral host-defense or viral pathogenesis, remain undefined.

In this study, we investigated the importance of necroptosis to human-pathogenic coronaviruses using *in vitro*, *ex vivo* human lung explant, and *in vivo* infection models. Our results show that necroptosis is most dramatically triggered by MERS-CoV, followed in degree by SARS-CoV-1 and SARS-CoV-2. In contrast, necroptosis activation is not detected in cells infected with seasonal coronaviruses, such as HCoV-NL63. We demonstrate that coronaviruses trigger necroptosis through the Z-DNA-binding protein 1 (ZBP1)-receptor-interacting serine/threonine-protein kinase 1 (RIPK1)/RIPK3-mixed lineage kinase domain like pseudokinase (MLKL) pathway. We further show that MERS-CoV infection induces rampant Z-RNA production, followed by SARS-CoV-1 and SARS-CoV-2, while little to no Z-RNA was detected in seasonal coronavirus (HCoV-229E, HCoV-OC43, HCoV-NL63)-infected cells, which explains the differential necroptosis activation upon coronavirus infection. Importantly, necroptosis inhibition by small-molecule based blockade of RIPK3 kinase activity markedly attenuated inflammatory cytokine expression, ameliorated lung damage, and rescued the survival of MERS-CoV-infected human DPP4 knock-in (hDPP4-KI) mice, without intervening virus clearance. Similarly, necroptosis blockade significantly reduced inflammation and lung pathology in K18-hACE2 mice infected with SARS-CoV-1 or SARS-CoV-2. Overall, our study pinpoints necroptosis as a key pathogenic driver of disease severity following infection by human-pathogenic coronaviruses, and positions small-molecule necroptosis inhibitors as a therapeutic strategy for all known highly-pathogenic coronaviruses, as well as for future zoonotic coronaviruses.

## RESULTS

### Necroptosis drives MERS-CoV-induced inflammation and cell damage

While necroptosis primarily aims to restrict virus replication, excessive necroptosis activation may result in heightened damage-associated molecular pattern (DAMP) release that exaggerate inflammatory response and tissue injury(*36, 37*). To explore if necroptosis is triggered in severe coronavirus infections, we utilized MERS-CoV as a model virus and investigated MERS-CoV infection in human *ex vivo* lung tissues (**Fig. 1A**). In the infected lung tissues, we detected abundant expression of phosphorylated MLKL (pMLKL), which is a hallmark of necroptosis (**Fig. 1B-C**). Interestingly, pMLKL was detected not only in infected nucleocapsid (N) protein positive cells (**Fig. 1B**, open arrowheads in inset i and ii), but was also present in N negative cells (**Fig. 1B**, dashed arrowheads in inset i and ii), indicating potential induction of necroptosis through a paracrine effect on bystander cells. As a control, pMLKL was not detected in mock-infected human *ex vivo* lung tissues (**Fig. 1B**). As MERS-CoV also triggers apoptosis in the infected human lung tissues(*12, 13*), we next delineated the relative contributions of apoptosis versus necroptosis to the MERS-CoV-induced pro-inflammatory response. To this end, we treated MERS-CoV-infected human lung tissues with the pan-caspase inhibitor, zVAD-FMK (zVAD), the MLKL membrane translocation inhibitor, necrosulfonamide (NSA), or a combination of both. Intriguingly, NSA but not zVAD treatment significantly attenuated the expression of inflammatory mediators, including key pro-inflammatory cytokines (TNF-α, CXCL10, IL-6, IL-1β) and inflammatory cell-recruiting chemokines (CXCL1, CXCL2, CXCL8, CCL2, CCL3) (**Fig. 1D and Fig. S1**), without reducing virus replication (**Fig. 1E**).

**Figure 1.**
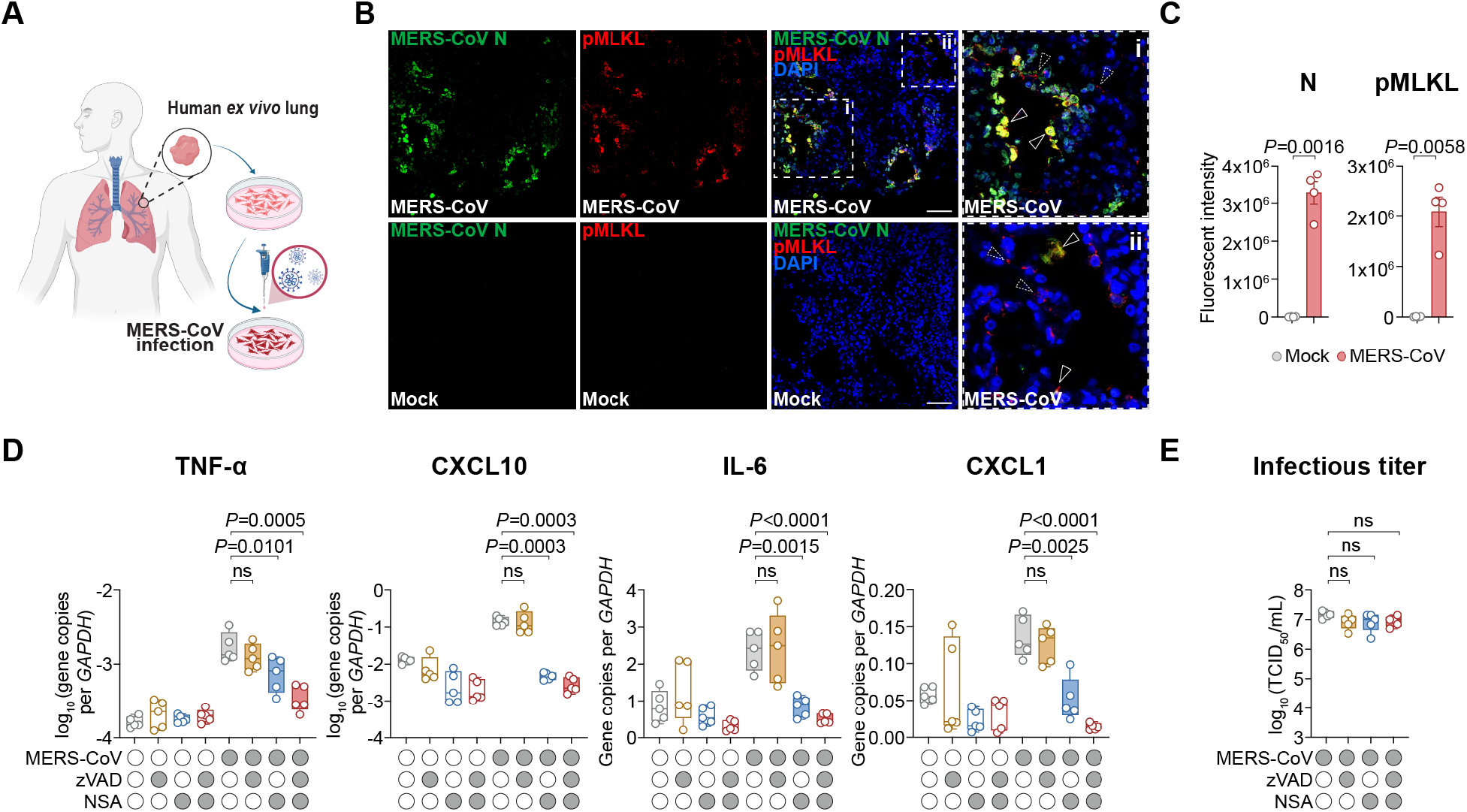
Necroptosis drives MERS-CoV-induced inflammation. (**A**) Human *ex vivo* lung tissues were inoculated with 3×10^6^ PFU MERS-CoV in 1 mL culture medium and harvested at 24 hpi. (**B**) MERS-CoV N and pMLKL expression was identified with immunostaining and (**C**) quantified with ImageJ (n=4). Mock-infected human *ex vivo* lung tissues were included as controls. (**D-E)** MERS-CoV-or mock-infected human *ex vivo* lung tissues were treated with 50 μM zVAD, 5 μM NSA, or a combination of both. Tissue lysate and supernatant samples were harvested at 24 hpi to determine the expression of pro-inflammatory markers (TNF-α, CXCL10, IL-6) and inflammatory cell-recruiting chemoattractant (CXCL1) (D), and viral titer (n=5) (E). Data represented mean and standard error of the mean (C) or median and interquartile range (D,E) from the indicated number of biological repeats. Statistical significance between groups was determined with two-tailed Student’s t-test (C) or one way-ANOVA with Tukey’s multiple comparisons tests (D,E). Bars in (B) represented 50 μm.

Next, we sought to delineate the molecular mechanism of MERS-CoV-induced necroptosis. Conventional cell lines commonly used for human-pathogenic coronavirus research including A549, BEAS2B, Calu3, Huh7, Caco2, 293T, and VeroE6, are deficient in necroptosis (**Fig. S2A**), likely due to defects in RIPK3 expression(*38, 39*). To this end, we utilized HT29 cells, which are widely used to assess virus-induced necroptosis (**Fig. S2A-B**). Necroptosis is mediated by RIPK1, RIPK3, and MLKL, and can be initiated when ZBP1 senses Z-RNA or Z-DNA triggered during virus infections (**Fig. 2A**). Our results demonstrated that MERS-CoV infection in HT29 cells dramatically upregulated the expression of ZBP1 and RIPK1 (**Fig. S3A**), and induced RIPK1, RIPK3, and MLKL phosphorylation in addition to caspase-3 cleavage in a dose and time dependent manner (**Fig. 2B and Fig. S3B**). In agreement with these observations, while zVAD or NSA treatment only modestly increased cell viability when applied alone, the combined treatment of zVAD and NSA more significantly rescued the survival of MERS-CoV-infected HT29 cells (**Fig. 2C-E**). Similar inhibition of MERS-CoV-induced cell death by zVAD and NSA was also detected in human intestinal organoids (**Fig. S3C-D**). To further dissect the contribution of RIPK1 and RIPK3 in MERS-CoV-induced necroptosis, we treated infected cells with RIPK1 inhibitor (necrostatin-1, Nec-1) or RIPK3 inhibitor (GSK872 or GSK840), in the presence of zVAD. We found that the combined treatment of zVAD/Nec-1, zVAD/GSK872, or zVAD/GSK840, all significantly increased cell viability (**Fig. 2D-E**). Taken together, these results indicate that MERS-CoV, like influenza viruses(*25, 40*), activates both necroptosis and apoptosis in infected cells, and that RIPK1 and RIPK3 are both required in MERS-CoV-induced necroptosis.

**Figure 2.**
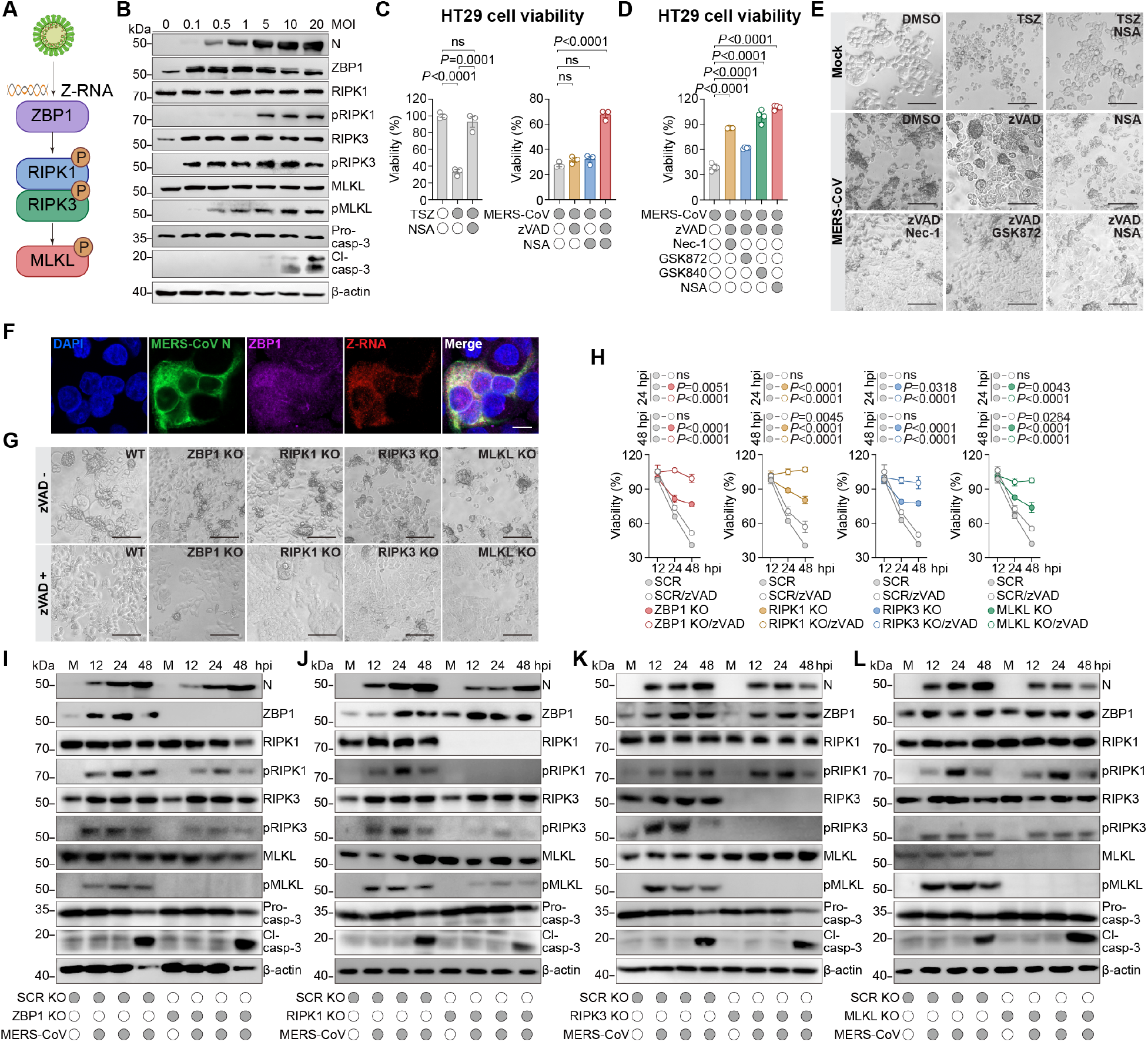
ZBP1 mediates MERS-CoV-induced necroptosis. (**A**) Schematic of the canonical ZBP1-mediated necroptosis pathway. (**B**) HT29 cells were infected with MERS-CoV at the indicated MOI and harvested at 24 hpi for Western blot analysis. (**C-E**) HT29 cells were infected with MERS-CoV at 10 MOI and treated with 5 μM NSA, 25 μM Nec-1, 3 μM GSK840, 5 μM GSK872, with or without 50 μM zVAD. Cell viability was quantified at 24 hpi with CellTiterGlo assays. 30 ng/mL TNF-α, 1 μM smac mimetic (BV6), and 25 μM zVAD (TSZ)-treated and TSZ/NSA-treated HT29 cells served as controls (n=3 for C and n=4 for D) (C,D). Cell morphology of the treated cells at 24 hpi (E). (**F**) MERS-CoV-infected HT29 was fixed at 24 hpi. The cellular localization of ZBP1 (purple), MERS-CoV N (green), and Z-RNA (red) was identified with immunofluorescence staining and confocal microscopy. (**G-L**) ZBP1, RIPK1, RIPK3, MLKL KO HT29 cells were infected with MERS-CoV at 10 MOI with or without 50 μM zVAD treatment. Cell morphology was imaged at 24 hpi (G). Cell viability was quantified at 12, 24, and 48 hpi with CellTiterGlo assays (n=4) (H). The expression of key components along the necroptosis and apoptosis pathway at 12, 24, and 48 hpi was detected with Western blots (I-L). Data represented mean and standard error of the mean from the indicated number of biological repeats. Statistical significance between groups was determined with one way-ANOVA with Tukey’s multiple comparisons tests (C,D) or two-way ANOVA with Šídák’s multiple comparisons test (H). Bars in (E, F, G) represented 100 μm, 10 μm, and 100 μm, respectively.

We next performed immunofluorescence staining assays and found that ZBP1 and its ligand Z-RNA were strongly colocalized in the MERS-CoV-infected cells (**Fig. 2F**). In particular, ZBP1 expression is upregulated in the infected cells (**Fig. 2F**), which is in keeping with our earlier observations from qPCR (**Fig. S3A**) and immunoblot analyses (**Fig. 2B**). To differentiate between Z-DNA and Z-RNA in the MERS-CoV-infected cells, we treated the infected cells with combinations of DNase I, RNase A, or RNase III. We found that while DNase I treatment of the infected cells did not reduce the Z-RNA immunostaining signal, combined RNase A and RNase III treatment reduced the Z-RNA signal to background level (**Fig. S3E**), indicating MERS-CoV infection induced Z-RNA but not Z-DNA production in the infected cells. Time course analysis of MERS-CoV-infected HT29 cells further revealed that Z-RNA production was readily detected at 12 hpi, before substantial pMLKL expression was detected. At 24 to 72 hpi, a high degree of Z-RNA and pMLKL co-expression was consistently detected in over 80% of the infected cells (**Fig. S3F-G**).

To establish a causal relationship between ZBP1 sensing and necroptosis induction upon MERS-CoV infection, we constructed ZBP1 knockout (KO) HT29 cells and investigated MERS-CoV-induced necroptosis in the ZBP1 KO cells. Our results revealed that ZBP1 KO completely abolished pMLKL expression and partially rescued MERS-CoV-induced cell death in the infected cells (**Fig. 2G-I**). Importantly, ZBP1 KO did not reduce caspase-3 cleavage (**Fig. 2I**), indicating that death pathways besides ZBP1 can also mediate apoptosis activation in MERS-CoV-infected cells. Notably, zVAD treatment in the ZBP1 KO cells completely rescued the viability of MERS-CoV-infected cells (**Fig. 2H**). Similarly, we constructed RIPK1, RIPK3, and MLKL KO HT29 cells. Upon infection, MERS-CoV triggered attenuated pMLKL expression and caspase-3 cleavage in the RIPK1 KO cells (**Fig. 2J**). In contrast, RIPK3 KO abolished pMLKL expression but did not impact caspase-3 cleavage (**Fig. 2K**). Interestingly, we observed a small but consistent increase in caspase-3 cleavage in the MLKL KO cells (**Fig. 2L**). In keeping with the ZBP1 KO results, RIPK1, RIPK3, and MLKL KO in part reduced MERS-CoV-induced cell death, and additional zVAD treatment consistently rescued the viability of all evaluated KO cells to nearly 100% (**Fig. 2G-H**). In parallel, we assessed MERS-CoV replication in the KO cells, and found that ZBP1, RIPK1, RIPK3, or MLKL KO did not enhance virus replication (**Fig. S3H**). Overall, these results demonstrate that ZBP1 senses coronavirus-generated Z-RNAs to initiate necroptosis, and that such necroptosis drives cell death and pro-inflammatory responses.

### Human-pathogenic coronaviruses differentially activate necroptosis

Next, to evaluate if necroptosis is triggered by other human-pathogenic coronaviruses, we infected HT29 cells modified to stably express human angiotensin-converting enzyme 2 (ACE2) (HT29-ACE2) (**Fig. S4A**) with SARS-CoV-1 and SARS-CoV-2 wildtype (WT), and assessed their potential of necroptosis activation, in comparison to that of MERS-CoV. Our results demonstrated that all three evaluated viruses robustly replicated in HT29-ACE2 cells (**Fig. 3A**) with concomitant induction of MLKL phosphorylation (**Fig. 3B**). Interestingly, at the same dose of inoculum (MOI=1), MERS-CoV induced significantly more pMLKL than that of SARS-CoV-1 and SARS-CoV-2 WT (**Fig. 3B**). Since these higher pMLKL levels may potentially result from the more efficient replication kinetics of MERS-CoV, compared to the other two viruses at the same MOI (**Fig. 3A and Fig. S4B**), we additionally challenged HT29-ACE2 cells with SARS-CoV-1 and SARS-CoV-2 WT with five times more infectious particles (5 MOI), which resulted in largely comparable levels of viral replication among the three viruses (**Fig. 3C and Fig. S4C**). Under these conditions, SARS-CoV-1 and SARS-CoV-2 WT did indeed trigger more pMLKL, but the levels of pMLKL remained substantially lower than that induced by MERS-CoV, and also manifested with notably slower kinetics (**Fig. 3D**). Interestingly, between the two Sarbecoviruses, SARS-CoV-1 induced more pMLKL production than did SARS-CoV-2 WT, despite replicating less efficiently than SARS-CoV-2 WT, consistent with the more severe disease triggered by SARS-CoV-1 (**Fig. 3C-D and Fig. S4C**). Of particular note, pMLKL was also readily detected in HT29-ACE2 cells infected with more recent SARS-CoV-2 variants (Omicron BA.1 and BA.5) (**Fig. S4D**).

**Figure 3.**
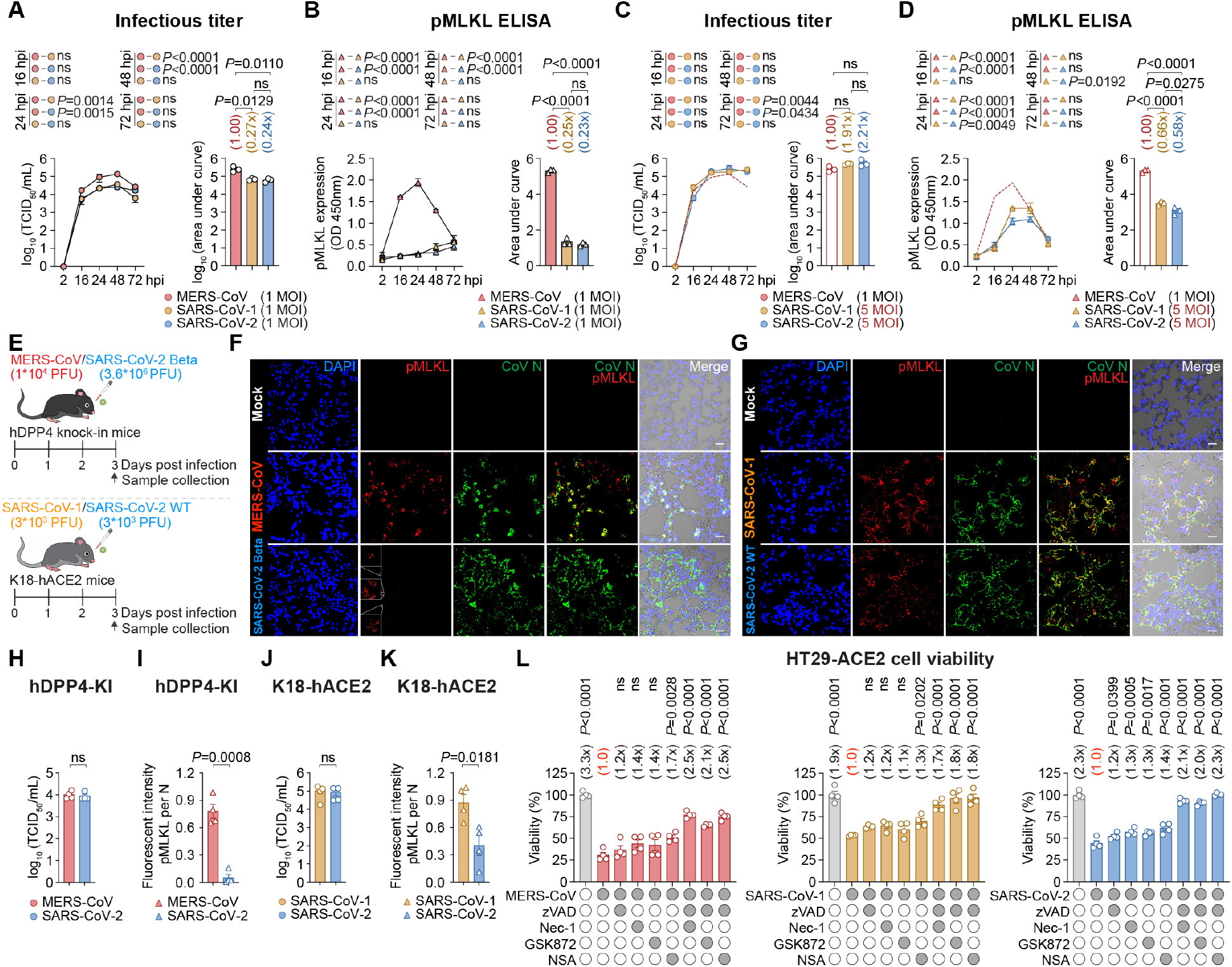
Human-pathogenic coronaviruses activate necroptosis in a pathogenicity-dependent manner. (**A-B**) HT29-ACE2 cells were infected with MERS-CoV, SARS-CoV-1, or SARS-CoV-2 wildtype (WT) at 1 MOI. Virus replication was determined with TCID_50_ assays using supernatant samples harvested at the indicated time points (n=3) (A). The level of pMLKL activation was quantified with ELISA using cell lysate samples harvested at the indicated time points (n=3) (B). (**C-D**) HT29-ACE2 cells were infected with SARS-CoV-1 or SARS-CoV-2 WT at 5 MOI. Virus replication (C) and pMLKL activation (D) was quantified with TCID_50_ assays and ELISA, respectively (n=3). (**E-K**) Schematic of *in vivo* pMLKL comparison. Coronavirus N and pMLKL expression in lung tissues of hDPP4-KI mice at 3 dpi were revealed with immunostaining (F). Coronavirus N and pMLKL expression in lung tissues of K18-hACE2 mice at 3 dpi were revealed with immunostaining (G). Virus replication of coronaviruses in hDPP4-KI and K18-hACE2 mice were assessed with TCID_50_ assays (n=4) (H,J). The pMLKL per N signal was quantified with ImageJ (n=4) (I,K). (**L**) HT29-ACE2 cells were infected with 10 MOI MERS-CoV, SARS-CoV-1, or SARS-CoV-2. The infected cells were treated with 25 μM Nec-1, 5 μM GSK872, 5 μM NSA, with or without 50 μM zVAD, and cell viability was quantified at 24 hpi with CellTiterGlo assays (n=4). Data represented mean and standard error of the mean from the indicated number of biological repeats. Statistical significance between groups was determined with two-tailed Student’s t-tests (H-K), one way-ANOVA with Tukey’s multiple comparisons tests (for the AUC analyses) and two-way ANOVA with Tukey’s multiple comparisons tests (for the replication curve analyses) (A-D, L). The same virus replication and pMLKL activation curve for MERS-CoV was used in (A,C) and (B,D), respectively. Bars in (F,G) represented 20 μm.

To examine if necroptosis is activated *in vivo* by the three highly-pathogenic coronaviruses, we examined pMLKL levels in infected mice at day 3 post virus challenge (**Fig. 3E-K**). By adjusting the inoculum of MERS-CoV and SARS-CoV-2 Beta, we achieved comparable levels of N gene expression (**Fig. S4E**), N protein expression (**Fig. 3F**), and infectious titer (**Fig. 3H**) between the two viruses in the hDPP4-KI mice infection model. Despite this equivalence in virus replication, we observed significantly more pMLKL in MERS-CoV-infected lung tissues when compared with that of SARS-CoV-2 Beta at 3 dpi (**Fig. 3F,I**). In parallel, we challenged K18-hACE2 mice with 3×10^3^ PFU/mice SARS-CoV-1 and SARS-CoV-2 WT, which resulted in comparable N gene expression (**Fig. S4F**), N protein expression (**Fig. 3G**), and infectious titer (**Fig. 3J**). In keeping with our observation in HT29-ACE2 cells, SARS-CoV-1 triggered significantly more robust pMLKL expression than that of SARS-CoV-2 WT in the lung tissues of infected mice at 3 dpi (**Fig. 3G,K**).

To delineate the mechanism of SARS-CoV-1- and SARS-CoV-2-induced necroptosis, we treated SARS-CoV-1- and SARS-CoV-2-infected HT29-ACE2 cells with Nec-1, GSK872, or NSA, with or without additional zVAD. Similar to the MERS-CoV control, SARS-CoV-1- and SARS-CoV-2-induced cell death was consistently rescued by the combined treatment of zVAD/Nec-1, zVAD/GSK872, or zVAD/NSA (**Fig. 3L**), indicating all three highly pathogenic coronaviruses may trigger necroptosis through a common pathway. HCoV-NL63 is a seasonal coronavirus that uses the same ACE2 host receptor for viral entry as SARS-CoV-1 and SARS-CoV-2(*41*). We infected HT29-ACE2 with HCoV-NL63 at 10 MOI and monitored pMLKL expression over a time course from 12 to 72 hpi. Strikingly, although the percentage of HCoV-NL63 N positive cells increased from 0.2% at 12 hpi to 35.6% at 72 hpi, pMLKL expression remained at baseline in the infected cells (**Fig. S4G-H**). As a control, 94.1% of N positive MERS- CoV-infected cells were pMLKL positive at 72 hpi (**Fig. S4G-H**). Together, our results demonstrate a striking correlation between necroptosis induction and coronavirus pathogenicity. MERS-CoV (35% mortality rate) most robustly triggered necroptosis, followed by SARS-CoV-1 (10% mortality rate), and then SARS-CoV-2 (1% mortality rate), while necroptosis was not detected in HCoV-NL63 (very low mortality)-infected cells.

We next sought to understand the mechanisms underlying the differential necroptosis induction between human-pathogenic coronaviruses. As Z-RNA is the upstream ligand sensed by ZBP1 for necroptosis initiation(*26, 42*) and as ZBP1 ablation abolished coronavirus-induced necroptosis (**Fig. 2I**), we investigated the propensity for Z-RNA formation by the different coronaviruses (**Fig. 4A-G**). In infected HT29-ACE2 cells, MERS-CoV triggered substantial and readily detectable Z-RNA formation (**Fig. 4A,E**). In comparison, SARS-CoV-1 and SARS-CoV-2 consistently induced Z-RNA formation in the infected cells but at a significantly lower level when compared to MERS-CoV, even after normalizing for N expression (**Fig. 4A,E**). Interestingly, no specific signal above background (mock-infection) level was detected in HCoV-NL63-infected cells (**Fig. 4A,E**). To confirm the lack of Z-RNA induction in HCoV-NL63-infected cells, we further examined HCoV-NL63 infection in LLC-MK2 cells, which is a model cell line for HCoV-NL63 studies that supports robust HCoV-NL63 replication (**Fig. S5A-B**). As a control, MERS-CoV triggered abundant Z-RNA expression in the infected LLC-MK2 cells at 24 hpi (**Fig. 4B**). Nevertheless, we were not able to detect any specific Z-RNA signal in HCoV-NL63-infected LLC-MK2 cells (**Fig. 4B**), in keeping with our findings in HT29-ACE2 cells.

**Figure 4.**
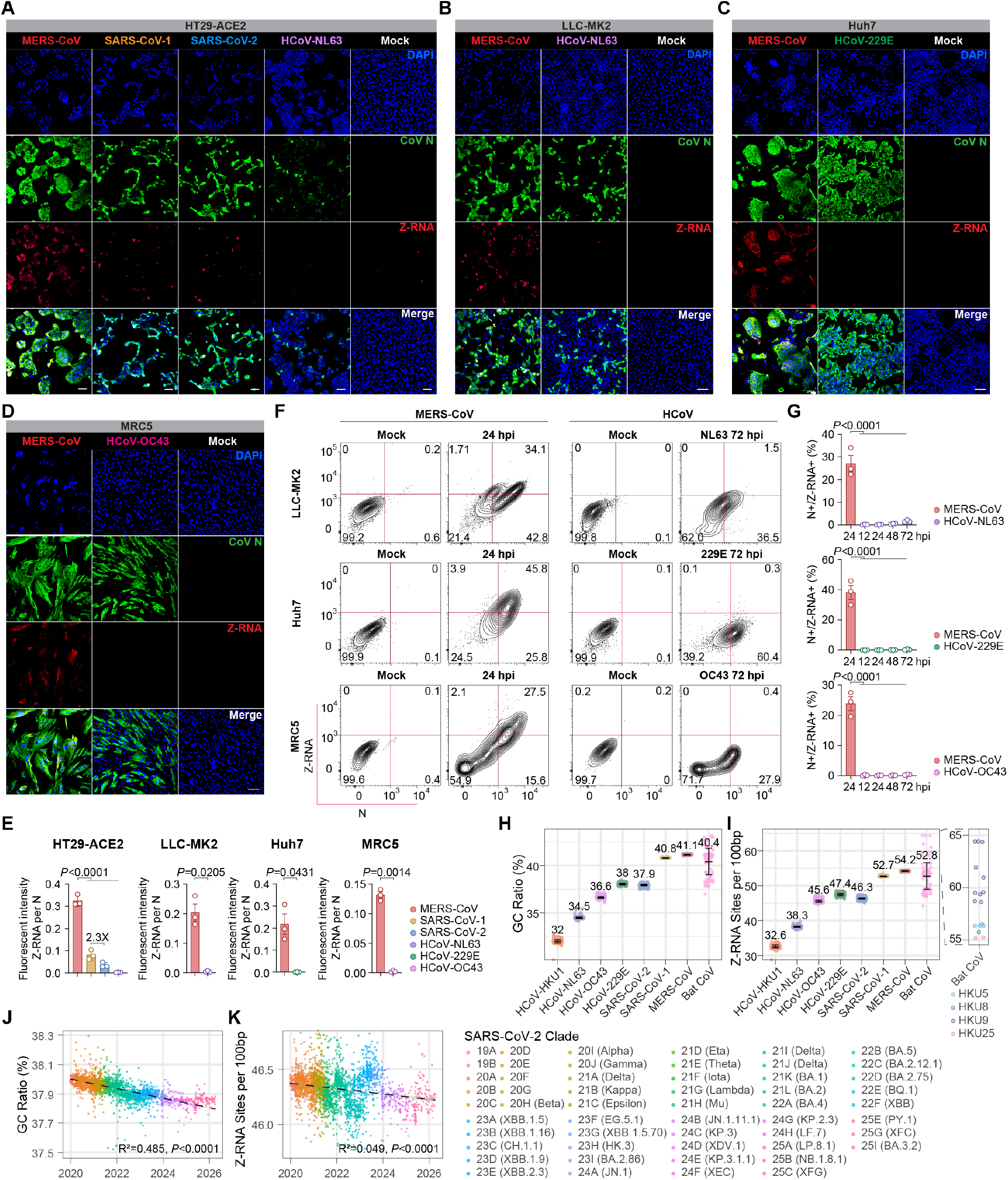
Z-RNA formation underlies differential human-pathogenic coronaviruses-triggered necroptosis. (**A-E**) Coronavirus-infected cells were fixed. Coronavirus N (green) and Z-RNA (red) expression were revealed with immunofluorescence staining and confocal microscopy. Cell nuclei were revealed with DAPI stain (blue). HT29-ACE2 cells were infected with MERS-CoV (1 MOI), SARS-CoV-1 (5 MOI), SARS-CoV-2 WT (5 MOI), or HCoV-NL63 (10 MOI), and harvested at 24 hpi (A). LLC-MK2 cells were infected with MERS-CoV (2 MOI) or HCoV-NL63 (1 MOI), and harvested at 72 hpi (B). Huh7 cells were infected with MERS-CoV (0.1 MOI) or HCoV-229E (10 MOI), and harvested at 24 hpi (C). MRC5 cells were infected with MERS-CoV (1 MOI) or HCoV-OC43 (10 MOI), and harvested at 24 hpi (D). Quantification of Z-RNA per coronavirus N signal from panel A-D with ImageJ (n=3) (E). (**F-G**) MERS-CoV- and seasonal coronavirus-infected cells were fixed at the indicated time points for detection of Z-RNA and N protein with flow cytometry. 5 MOI of MERS-CoV and seasonal coronaviruses were used for each cell type (F). Summary of the Z-RNA and N double positive cells at the indicated time points (n=3) (G). (**H**) GC content ratio of human-pathogenic and bat coronaviruses. (**I**) Z-score of human-pathogenic and bat coronaviruses predicted by ZHUNT3. Bat coronaviruses with the highest predicted Z-scores were identified. (**J-K**) GC content ratio and Z-score of SARS-CoV-2 variants predicted by ZHUNT3. 2,885 SARS-CoV-2 sequences collected from GenBank between 2020 and 2026 were categorized by clade to capture the evolutionary trajectory. Data represented mean and standard error of the mean from the indicated number of biological repeats. Statistical significance between groups was determined with two-tailed Student’s t-test (E), one way-ANOVA with Tukey’s multiple comparisons tests (E,G), or linear regression (J,K). Bars in (A) and in (B-D) represented 200 μm and 100 μm, respectively.

The absence of Z-RNA in HCoV-NL63-infected cells prompted us to assess whether Z-RNA is generated by other seasonal coronaviruses. To this end, we evaluated HCoV-229E and HCoV-OC43 infection in Huh7 and MRC5 cells, respectively, with MERS-CoV included as controls. We first infected Huh7 cells with MERS-CoV and HCoV-229E in parallel. At 24 hpi, abundant MERS-CoV and HCoV-229E N protein was identified in the infected cells, indicating robust replication of both viruses. However, we were only able to detect Z-RNA in MERS-CoV-infected cells but not HCoV-229E-infected cells (**Fig. 4C,E**). Similarly, in MRC5 cells infected with MERS-CoV and HCoV-OC43, Z-RNA was detected following MERS-CoV infection, but not after HCoV-OC43 infection (**Fig. 4D-E**). To further evaluate Z-RNA formation in seasonal coronavirus-infected cells in a more quantitative manner, we performed flow cytometry-based time course assays to measure Z-RNA production. At 24 hpi, 44.3% (34.1/76.9) of N positive MERS-CoV-infected LLC-MK2 cells were Z-RNA positive, while the proportion of Z-RNA positive cells among N positive MERS-CoV-infected Huh7 and MRC5 cells were 64.0% (45.8/71.6) and 63.8% (27.5/43.1), respectively (**Fig. 4F-G**). In contrast, no Z-RNA was found in seasonal coronavirus-infected cells at 24 hpi (**Fig. S5C**). We then extended our analysis to longer time points including 72 hpi for seasonal coronavirus-infected cells. Under this setting, we were able to identify a small population of Z-RNA positive cells within the N positive HCoV-NL63-infected LLC-MK2 cells (1.5/38.0; 3.9%). However, no Z-RNA signal in HCoV-229E-and HCoV-OC43-infected cells was identified (**Fig. 4F-G and Fig. S5C**). We subsequently examined HCoV-229E- and HCoV-OC43-infected cells with immunofluorescence staining and confocal microscopy from 24 to 72 hpi, and no Z-RNA signal in these cells was detected despite the development of virus-induced cytopathic effects at the late time points. In contrast, Z-RNA was detected in the same set of cells upon MERS-CoV infection at as early as 6 hpi (**Fig. S5D-E**). Taken together, these results indicate differential Z-RNA induction may underlie the difference in necroptosis activation between human-pathogenic coronaviruses.

Throughout our investigation, MERS-CoV consistently induced the highest level of Z- RNA in the infected cells. Since the propensity for viral Z-RNA formation is strongly correlated with the guanine and cytosine (GC) content of the nucleotide sequence(*27*), we analyzed the GC contents of MERS-CoV and other coronavirus genomes, and calculated the probability of Z-formation (the Z-score) using the ZHUNT3 algorithm(*43, 44*). Our analysis revealed that MERS-CoV has the highest genomic GC content ratio and Z-score among all human-pathogenic coronaviruses (**Fig. 4H-I**). Notably, MERS-CoV and SARS-CoV-1, both of which have higher genomic GC content and Z-scores than any of the other human-pathogenic coronaviruses, have only limited circulation in humans (**Fig. 4H-I**). Since both MERS-CoV and SARS-CoV-1 are zoonotic viruses that originate from bats(*2, 7*), we investigated the GC content ratio and Z-score of 111 distinct bat coronaviruses. Interestingly, we found that bat coronaviruses are in general high in GC content ratio and Z-scores, similar to that of MERS-CoV and SARS-CoV-1 (**Fig. 4H,I**). Among the bat coronaviruses, HKU5, HKU8, HKU9, and HKU25 were found to have the highest Z-score, even higher than that of MERS-CoV (**Fig. 4I**). Given that SARS-CoV-2 has now become endemic in humans(*45, 46*). We probed for the GC content ratio and Z-scores of SARS-CoV-2 isolates from 2020, when it was a recent zoonotic entrant into human populations, to the present time (2026), when it has started to adapt to humans. Our results demonstrate a clear decrease in GC content ratio for SARS-CoV-2 isolates over time (**Fig. 4J**). In agreement with these observations, a remarkably significant (p<0.0001) decrease in Z-score was observed as SARS-CoV-2 has adapted to humans over the past six years (**Fig. 4K**). Overall, these results are consistent with the idea that highly pathogenic (and recently zoonotic) coronaviruses generate greater levels of Z-RNA and activate robust ZBP1-dependent necroptosis, but subsequent human adaptation likely selects for progeny viruses with lower propensity to form Z-RNA, trigger ZBP1, and induce deleterious inflammation.

### Necroptosis inhibition attenuates severe coronavirus infections

Exaggerated inflammatory response and lung injury are often observed in patients infected by highly pathogenic coronaviruses but whether necroptosis plays a role in driving pathology during severe coronavirus infections is unknown. To investigate if necroptosis inhibition attenuates severe coronavirus pathogenesis, we intranasally inoculated human DPP4 knock-in (hDPP4-KI) mice with mouse-adapted MERS-CoV (MERS-CoV_MA_)(*47*) (**Fig. 5A**). The virus-challenged mice were then treated with the pan-caspase inhibitor zVAD, the RIPK3 kinase inhibitor GSK872, or both agents, for six consecutive days and monitored for bodyweight change and survival until 14 dpi (**Fig. 5B-C**). In parallel, a subset of challenged animals was examined on 3 dpi for virological and pathological analyses (**Fig. 5D-H**). We found that GSK872 treatment alone increased animal survival from ∼36% to >80% (*P* = 0.0205), while zVAD treatment alone did not improve animal survival (**Fig. 5B-C**). Combined zVAD and GSK872 treatment also improved body weight and increased animal survival, but it was not significantly greater than the protection offered by GSK872 alone (**Fig. 5B-C**). In the virological assessments, we found that none of the single or combined treatment significantly altered virus replication (**Fig. 5D**). Nevertheless, GSK872 and the combined zVAD/GSK872 treatment significantly reduced the expression of representative pro-inflammatory cytokines, TNF-α and CXCL10 (**Fig. 5E**), and markedly decreased pMLKL expression (**Fig. 5F-G**) in the lung tissue of infected animals. In agreement with these findings, histological assessment of infected lung tissues indicated that inhibition of necroptosis markedly ameliorated MERS-CoV-induced lung damage. While severe bronchiolar epithelial cell death and desquamation, extensive mononuclear cell infiltration of alveolar space, massive protein-rich fluid exudation and haemorrhage, and focal alveolar wall collapse were found in the lungs of vehicle-treated mice at 3 dpi, GSK872 or the combined zVAD/GSK872 treatment significantly alleviated these histopathological changes (**Fig. 5H-I**). Interestingly, we observed more alveolar exudation, haemorrhage, and more severe destruction of alveolar structure in the zVAD-treated mice than that in the vehicle-treated mice (**Fig. 5H-I**), which corroborated with the higher pMLKL expression (**Fig. 5F-G**), more significant body weight drop (**Fig. 5B**), and earlier onset of animal death (**Fig. 5C)**, likely due to zVAD-mediated caspase-8 inhibition and consequent increased necroinflammatory injury.

**Figure 5.**
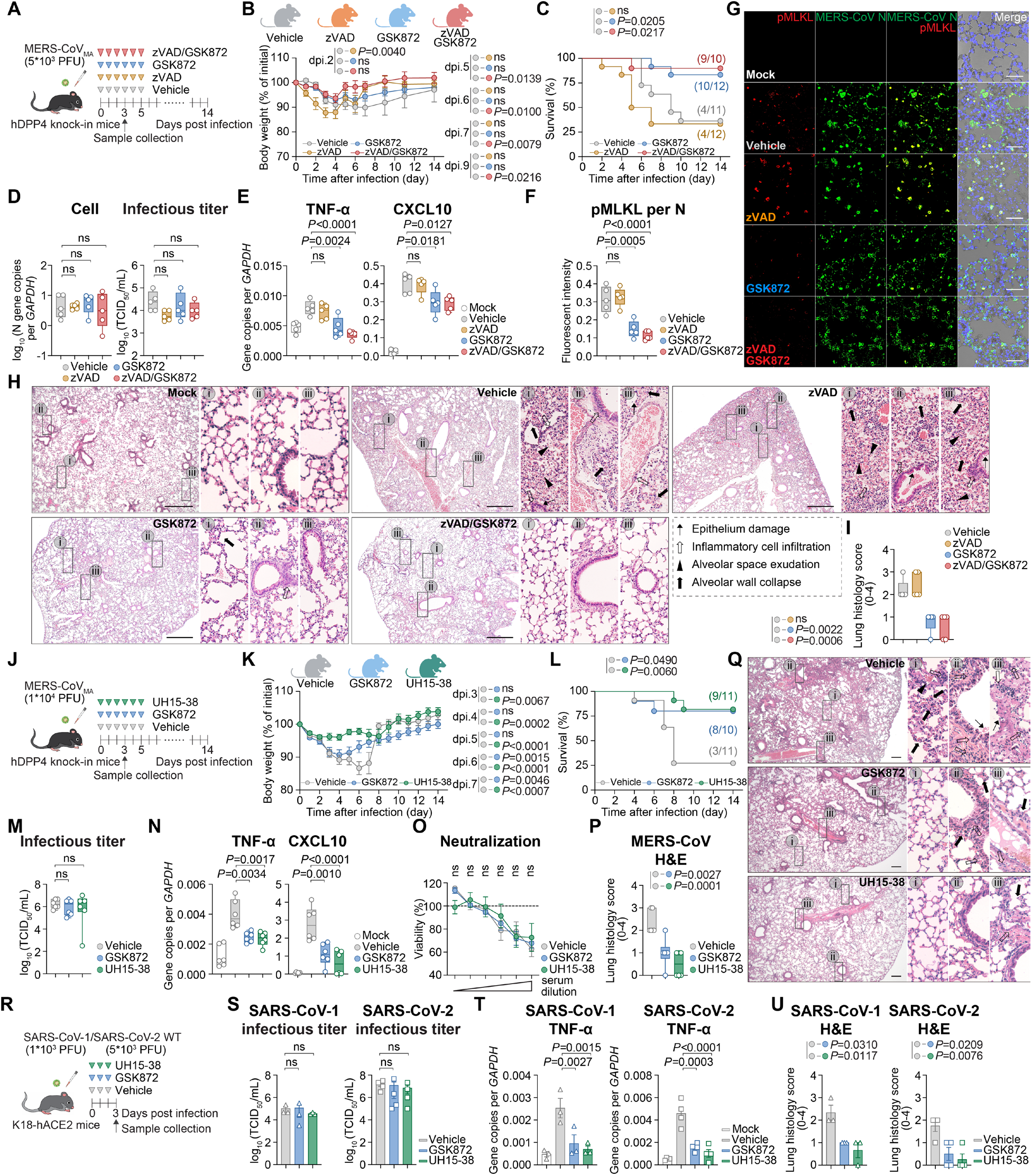
Necroptosis inhibition prevents lung injury in severe coronavirus infection. (**A**) Schematic of animal experiment with zVAD and GSK872 treatment. hDPP4-KI mice were inoculated with 5×10^3^ PFU/mouse MERS-CoV_MA_ through the intranasal (i.n.) route followed by intraperitoneal (i.p.) administration of 12.5 mg/kg/day zVAD, 10 mg/kg/day GSK872, or a combination of both, or vehicle, for six consecutive days. (**B**) Bodyweight change (**C**) and survival were monitored until 14 dpi. A subset of animals was sacrificed at 3 dpi. (**D**) Virus N gene expression and infectious titer was determined with RT-qPCR and TCID_50_ assays, respectively (n=5). (**E**) Expression level of inflammatory cytokines, TNF-α and CXCL10, were quantified with RT-qPCR (n=5). (**F-G**) Viral N protein (green) and pMLKL (red) expression in lung tissues were revealed with immunofluorescence staining and quantified with ImageJ (n=5). (**H-I**) Representative images of histopathological changes in the lung revealed with hematoxylin and eosin (H&E) staining. Four categories of characteristic histopathological changes including epithelium damage, inflammatory cell infiltration, alveolar space exudation and alveolar wall collapse were quantified and scored (n=5). (**J**) Schematic of animal experiment with UH15-38 or GSK872 treatment. hDPP4-KI mice were inoculated with 1×10^4^ PFU/mouse MERS-CoV_MA_ through the i.n. route followed by i.p. administration of 30 mg/kg/day UH15-38, 10 mg/kg/day GSK872, or vehicle for six consecutive days. (**K**) Bodyweight change (**L**) and survival were monitored until 14 dpi. A subset of animals was sacrificed at 3 dpi. (**M**) Infectious titer was determined with TCID_50_ assays (n=6). (**N**) Expression level of inflammatory cytokines, TNF-α and CXCL10, were quantified with RT-qPCR (n=6). (**O**) Serum was harvested from survived animals on 16 dpi. *In vitro* neutralization assays were performed against MERS-CoV_MA_ (n=3 for vehicle, n=5 for GSK872 and UH15-38). (**P-Q**) Representative images of histopathological changes in the lung revealed with H&E staining. Four categories of characteristic histopathological changes including epithelium damage, inflammatory cell infiltration, alveolar space exudation and alveolar wall collapse were quantified and scored (n=6). (**R**) Schematic of animal experiment with UH15-38 or GSK872 treatment upon SARS-CoV-1 or SARS-CoV-2 infection. K18-hACE2 mice were inoculated with 1×10^3^ PFU/mouse SARS-CoV-1 or 5×10^3^ PFU/mouse SARS-CoV-2 WT through the i.n. route followed by i.p. administration of 30 mg/kg/day UH15-38, 10 mg/kg/day GSK872, or vehicle for three consecutive days. (**S**) Infectious titer was determined with TCID_50_ assays (n=3). (**T**) TNF-α expression was quantified with RT-qPCR (n=3). (**U**) Histopathological changes in the lung were revealed with H&E staining and scored (n=3). Data in (S-U) represented mean and standard error of the mean from the indicated number of biological repeats. Statistical significance between groups was determined with one way-ANOVA with Tukey’s multiple comparisons tests (D,E,F,I,M,N,P,S,T,U), Log-rank (Mantel-Cox) tests (C,L), or two way-ANOVA with Dunnett’s (B,K) or Šídák’s (O) multiple comparisons tests. Bars in (G,H,Q) represented 50 μm, 500 μm, and 200 μm, respectively.

Since necroptosis inhibition with GSK872 alone offered robust protection, we next sought to explore the efficiency of necroptosis inhibition on MERS-CoV pathogenesis. As GSK872 was reported to trigger on-target RIPK3-dependent apoptosis at high concentrations(*48*), we also utilized UH15-38, a well-tolerated and highly potent RIPK3 kinase inhibitor specifically developed for *in vivo* use(*49*), in these studies. We intranasally inoculated hDPP4-KI mice with a higher inoculum of MERS-CoV_MA_ (1×10^4^ PFU/mouse), treated the mice once-daily with either GSK872 or UH15-38 for six consecutive days after infection, and monitored for bodyweight change and survival until 14 dpi, while a subset of challenged animals was examined on 3 dpi for virological and pathological analyses (**Fig. 5J**). In this setting, we found that UH15-38 potently attenuated virus-induced body weight loss and increased mouse survival from ∼27% to >80% (P = 0.0060) (**Fig. 5K-L**). Similar to GSK872, UH15-38 did not alter virus replication kinetics but significantly reduced the expression of key pro-inflammatory cytokines, TNF-α and CXCL10, in the lung tissues of the infected animals (**Fig. 5M-N**). We further showed that necroptosis inhibition did not reduce neutralization antibody production, which was evidenced by the comparable neutralization capacity against MERS-CoV_MA_ using serum harvested from survived mice with GSK872, UH15-38, or vehicle treatment, at 16 dpi (**Fig. 5O**). Histological analysis revealed that UH15-38 robustly alleviated virus-induced lung injury. When compared with GSK872, UH15-38 was more effective at decreasing inflammatory cell infiltration and alveolar wall collapse, resulting in a lower pathology score (**Fig. 5P-Q**), which is in keeping with the significantly reduced body weight loss observed upon UH15-38 treatment (**Fig. 5K**).

We demonstrated earlier that all three highly pathogenic coronaviruses triggered necroptosis in the lung tissues of the infected animals (**Fig. 3F-G**). To further explore the potential link of necroptosis in COVID-19 severity, we analyzed single-cell RNA sequencing (scRNA-seq) data of bronchoalveolar lavage fluid (BALF) obtained from SARS-CoV-2 Omicron-infected or control subjects (**Fig. S6A-B**). We found that among the ciliated cells that are the primary targets of SARS-CoV-2 infection, markedly upregulated expression of ZBP1 was observed in the fatal cases when compared with the mild cases or uninfected cases (**Fig. S6C-D**). To investigate if necroptosis inhibition protects against SARS-CoV-1 and SARS-CoV-2 pathogenesis, we challenged K18-hACE2 mice with 1×10^3^ PFU/mouse SARS-CoV-1 or 5×10^3^ PFU/mouse SARS-CoV-2 WT, treated the mice with GSK872 or UH15-38 for three consecutive days, and sacrificed the infected mice for sample collection at 3 dpi (**Fig. 5R**). Consistent with our findings on MERS-CoV, GSK872 and UH15-38 treatment did not suppress the virus replication of SARS-CoV-1 and SARS-CoV-2 but efficiently reduced the expression of TNF-α in the lung tissues, and markedly ameliorated lung pathologies including epithelium damage, inflammatory cell infiltration, alveolar space exudation and alveolar wall collapse (**Fig. 5S-U and Fig. S7**). Together, these results demonstrate that necroptosis inhibition effectively reduces the extent of highly pathogenic coronavirus-triggered lung injury, in manner independent of virus clearance.

## DISCUSSION

In this study, we unambiguously show that necroptosis is a key and targetable instigator of pathogenesis during severe coronavirus infections. Using a spectrum of *in vitro*, *ex vivo*, and *in vivo* systems, we demonstrate that necroptosis is triggered by all three highly pathogenic coronaviruses (MERS-CoV, SARS-CoV-1, SARS-CoV-2). Inhibition of necroptosis in MERS-CoV-infected *ex vivo* human lung tissues significantly reduced the expression of pro-inflammatory cytokines and inflammatory chemoattractants. *In vivo*, necroptosis inhibition via small molecule RIPK3 kinase blockade significantly reduced body weight loss, attenuated virus-induced histopathological changes in the lungs, and improved the survival of MERS-CoV-infected animals. In keeping with our findings on MERS-CoV, necroptosis inhibition similarly reduced pro-inflammatory cytokine expression and significantly ameliorated lung injury of SARS-CoV-1-and SARS-CoV-2-infected K18-hACE2 mice. Most importantly, the attenuation of severe coronavirus-induced pathology was achieved independent of virus inhibition. This is in contrast to DNA virus infection in which necroptosis restricts viral replication(*50*). These results indicate that necroptosis inhibition does not suppress beneficial antiviral immune responses. Instead, it selectively dampens necroinflammatory lung injury, without impeding virus clearance. As future emergence of zoonotic coronaviruses remains high, the virus-independent nature of necroptosis targeting represents a potential broad-spectrum strategy to counter the threat of emerging highly pathogenic coronaviruses (**Fig. S8**).

Interestingly, we identified a positive correlation between the level of necroptosis activation and the degree of coronavirus pathogenicity. We found that MERS-CoV triggered the highest level of necroptosis, followed by SARS-CoV-1, and then by SARS-CoV-2. The level of necroptosis activation strongly aligns with the case-mortality rate of the three viruses at 35%, 10%, and 1%, respectively. Importantly, the comparisons were conducted using authentic viruses with *in vitro* and *in vivo* models properly normalized for differences in virus replication, which increases the credibility of these findings, and resolves existing controversy of the role of necroptosis in SARS-CoV-2 pathogenesis(*31–35*). In agreement with these results, scRNA-seq of BALF obtained from SARS-CoV-2 Omicron-infected or control subjects demonstrated that the ZBP1 necroptosis pathway is significantly more upregulated in the fatal cases when compared with the mild cases or uninfected cases. In our comparative analysis, we further demonstrated that the seasonal coronavirus HCoV-NL63 does not trigger necroptosis in the infected cells despite abundant N protein expression, while the positive control MERS-CoV triggered robust necroptosis under the same settings. In this regard, our study implicates necroptosis as a critical determinant that differentiates highly pathogenic coronaviruses from seasonal coronaviruses. While several other viral factors including receptor usage, receptor distribution, cell tropism, and immune evasion potential, are also known to regulate coronavirus pathogenicity(*51–57*), our study suggests that the propensity to activate ZBP1-driven necroptosis is a key determinant of coronavirus virulence.

By comparing the infection of authentic highly pathogenic coronaviruses (MERS-CoV, SARS-CoV-1, SARS-CoV-2) and the common cold related seasonal coronaviruses (HCoV-NL63, HCoV-229E, HCoV-OC43) in the same set of cells, we unexpectedly found that Z-RNA was readily detected in highly pathogenic coronavirus-infected cells but was mostly undetectable in seasonal coronavirus-infected cells. Among the highly pathogenic coronaviruses, MERS-CoV triggered the highest level of Z-RNA production, followed by SARS-CoV-1 and then SARS-CoV-2. This differential Z-RNA production may mechanistically explain why MERS-CoV activates the highest degree of necroptosis, and SARS-CoV-2 the least. Despite our efforts of using high MOI and detection at multiple time points, little to no Z-RNA production was detected in HCoV-NL63-, HCoV-229E-, or HCoV-OC43-infected cells. The underlying reasons for the low Z-RNA production by seasonal coronaviruses are not completely understood. It is possible that the original zoonotic hosts of coronaviruses including birds and bats may be defective or dampened in the necroptosis machinery, which allows Z-RNA-producing coronaviruses to co-evolve with the hosts. In support to this perspective, birds do not have ZBP1 or RIPK3. Bats, on the other hand, are reported to have an intact necroptosis machinery, but whether the ZBP1 pathway is fully functional in these mammals, has not been determined(*58, 59*). Given that inflammasome and other innate immune antiviral signaling are dampened in bats(*60–63*), it is possible that necroptosis signaling in bats may also be reduced, allowing them to tolerate coronaviruses and other potentially necroptogenic viruses, without deleterious consequences.

It is tempting to speculate that since humans have an intact necroptosis machinery, once coronaviruses jump into humans, human adaptation will select for viruses with low Z-RNA formation propensity. This speculation is supported by the finding that coronaviruses without human adaption (MERS-CoV, SARS-CoV-1) have significantly higher Z-scores than that of common cold coronaviruses (HCoV-NL63, HCoV-229E, HCoV-OC43, HCoV-HKU1) that have circulated in humans for years. In keeping with this hypothesis, our analysis has revealed a gradual but clearly observable and significant (p<0.0001) decrease in Z-scores for SARS-CoV-2 from 2020 to 2026. An exception to this trend is the ancestral SARS-CoV-2, whose Z-forming propensity is closer to the seasonal coronaviruses than it is to either MERS-CoV or SARS-CoV-1. Although we do not have a clear explanation for why this is the case, we speculate that ancestral SARS-CoV-2 may have been circulating in an intermediate host with an intact necroptosis machinery that has driven SARS-CoV-2 evolution towards low Z-RNA formation, prior to its jump to humans. Nevertheless, once in humans, SARS-CoV-2 genomes appear to be subject to additional selection pressure against Z-RNA formation and ZBP1 activation. Alternatively, host cell Z-RNA may contribute to ZBP1 activation by coronaviruses, as has recently been shown for influenza A virus (IAV) and herpes simplex virus 1 (HSV-1)(*64*). Both IAV and HSV-1 activate host cell Z-RNA production via disruption of transcription termination (DoTT)(*64*). In this regard, the ability to activate host cell Z-RNA production may be a determinant of coronavirus pathogenicity, and therefore an additional factor in driving selection of SARS-CoV-2 for human adaptation. This possibility may account for the lower Z-score of the ancestral SARS-CoV-2 genome, and remains to be experimentally examined. Nonetheless, viral genome-derived Z-RNA is likely important since it can adopt the left-handed Z-RNA conformation early in the infection cycle as part of the virus replication process, and serves to initiate the ZBP1 necroptosis cascade before massive host Z-RNA is induced by viral proteins. By analyzing the bat coronaviruses, we identified coronaviruses with high Z-scores, including the Rousettus bat coronavirus HKU9. HKU9 is a Nobecovirus (Betacoronavirus lineage D) first reported by us in 2006(*65*). We later demonstrated that HKU9 can utilize human 78-kDa glucose-regulated protein (GRP78) as an attachment factor(*66*), indicating its potential as a zoonotic pathogen. Therefore, surveillance programs should be in place for coronaviruses with high Z-score genomes (such as HKU9) as they may pose an especially imminent threat to humankind if they are able to jump the species barrier and infect humans.

Overall, our study positions necroptosis as a critical determinant of pathogenesis triggered by highly pathogenic coronaviruses, and establish necroptosis inhibition as an effective and readily-deployable strategy to attenuate the necroinflammatory pathogenesis of current and future virulent coronaviruses.

## MATERIALS AND METHODS

### Virus and safety

The MERS-CoV (GenBank: JX869059.2) was a kind gift from R. Fouchier (Erasmus Medical Center, Rotterdam, The Netherlands)(*67, 68*). The mouse-adapted MERS-CoV (MERS-CoV_MA_) was a kind gift from P. McCray (University of Iowa, IA, USA). HCoV-OC43 (GenBank: AY391777) was a kind gift from J. Zhao (Guangzhou Medical University, China) and X. Ma (Guangzhou National Laboratory, China). SARS-CoV-2 wild-type HKU-001a (GenBank: MT230904), Omicron BA.1 (GenBank: OM212472) and Omicron BA.5 (GISAID: EPI_ISL_13777658) were isolated from laboratory-confirmed patients with COVID-19 in Hong Kong(*69*). SARS-CoV-1 GZ50 (GenBank: AY304495), HCoV-229E (GenBank: PQ243243) and HCoV-NL63 were archived clinical isolates at Department of Microbiology, The University of Hong Kong. MERS-CoV and SARS-CoV-1 were propagated and titrated by plaque assays in VeroE6 cells. All variants of SARS-CoV-2 were cultured and titrated by plaque assays using VeroE6-TMPRSS2 cells. HCoV-229E was propagated and titrated by median tissue culture infectious dose (TCID_50_) assays in Huh7 cells. HCoV-OC43 and HCoV-NL63 were propagated in MRC5 and LLC-MK2 cells, respectively, and titrated by immunofluorescence staining. After obtaining the virus culture, the sequences of all variants used in this study were confirmed with nanopore sequencing. All infectious experiments involving MERS-CoV, SARS-CoV-2 and SARS-CoV-1 followed the approved standard operating procedures of the Biosafety Level 3 (BSL-3) facility at the Department of Microbiology, HKU.

### Animals and animal ethical approval

The human DPP4 exon 10 to 12 knock-in (hDPP4-KI) mice were provided by P. McCray (University of Iowa, IA, USA). Heterozygous K18-hACE2 C57BL/6J mice (2B6.Cg-Tg(K18-ACE2)2Prlmn/J) were obtained from The Jackson Laboratory. The animals were kept in cages with individual ventilation under 65% humidity and an ambient temperature of 21-23 °C and a 12-12 h day-night cycle for housing and husbandry. Food and water were provided to the animals without restriction. Group sizes were chosen based on statistical power analysis and our prior experience(*12, 13, 45, 46, 70*). Gender-and age-matched mice were randomized into different experimental groups. All animal experiments were approved by the Committee on the Use of Live Animals in Teaching and Research (CULATR) of The University of Hong Kong (HKU) under CULATR 24-313.

### Necroptosis activation in virus-infected mice

On the day of infection, 12-to 16-week-old hDPP4-KI mice anaesthetized with 100 mg/kg ketamine and 10 mg/kg xylazine were intranasally inoculated with 1×10^4^ PFU/mouse MERS-CoV or 3.6×10^6^ PFU/mouse SARS-CoV-2 Beta variant pre-diluted in 20 μl DMEM. 8-to 12-week-old K18-hACE2 transgenic mice were intranasally inoculated with 3×10^3^ PFU SARS-CoV-1 or SARS-CoV-2 WT pre-diluted in 20 μL DMEM. At 3 dpi, infected mice were sacrificed and lung tissues were harvested for virological assessment by RT-qPCR, TCID_50_ assays, and immunofluorescence staining. Lung tissues from mock-infected mice were included as controls.

### Virus challenge and drug treatment in mice

On the day of infection, 12-to 16-week-old hDPP4-KI mice or 8-to 12-week-old K18-hACE2 mice were anaesthetized with 100 mg/kg ketamine and 10 mg/kg xylazine. For pMLKL detection, the hDPP4-KI mice were intranasally inoculated with 1×10^4^ PFU/mouse MERS-CoV or 3.6×10^6^ PFU/mouse SARS-CoV-2 Beta, pre-diluted in 20 μL DMEM. The K18-hACE2 mice were intranasally inoculated with 3×10^3^ PFU/mouse SARS-CoV-1 and SARS-CoV-2 WT, pre-diluted in 20 μL DMEM. At 3 dpi, mice lung tissues were collected for RT-qPCR, TCID_50_ and immunofluorescence staining. For necroptosis inhibition, the hDPP4-KI mice were intranasally inoculated with 5×10^3^ or 1×10^4^ PFU MERS-CoV_MA_ pre-diluted in 20 μL DMEM, followed by intraperitoneal administration with 12.5 mg/kg zVAD-FMK, 10 mg/kg GSK872 (both dissolved in 400 μL 0.3% methylcellulose/0.1% tween-80/PBS), a combination of zVAD-FMK and GSK872, 30 mg/kg UH15-38 (dissolved in 400 μL 10% solutol HS-15/PBS) or vehicle only, once per day for 6 days or until sample collection. The health status and body weight of the mice were monitored for 14 days on a daily basis or until the mice were euthanized because of reaching the humane endpoint of the experiment. A subset of mice was euthanized at 3 dpi and lung tissues from mice of both treatment and vehicle groups were collected for RT-qPCR, TCID_50_, immunofluorescence staining, and H&E staining. For necroptosis inhibition, the K18-hACE2 mice were intranasally inoculated with 1×10^3^ PFU/mouse SARS-CoV-1, or 5×10^3^ PFU/mouse SARS-CoV-2 WT pre-diluted in 20 μL DMEM, followed by intraperitoneal administration with 10 mg/kg GSK872 (dissolved in 400 μL 0.3% methylcellulose/0.1% tween-80/PBS), 30 mg/kg UH15-38 (dissolved in 400 μL 10% solutol HS-15/PBS) or vehicle only, once per day for 3 days. At 3 dpi, mice lung tissues were collected for RT-qPCR, TCID_50_, immunofluorescence staining, and H&E staining.

### Serum neutralization assay

Neutralization assays were performed using serum collected from hDPP4 knock-in mice at 16 days post-infection. Sera from the vehicle, UH15-38, and GSK872 treated groups were serially diluted in 5-fold dilutions and incubated with 100 PFU of MERS-CoV_MA_ for 1 h at 37 °C before adding to VeroE6 cells seeded in 96-well plates. Each dilution was tested in quintuplicate. Cells were maintained for 5 days at 37 °C in a humidified incubator with 5% CO_2_. At 5 days post-infection, cytopathic effect (CPE) was assessed by visual inspection. In parallel, cell viability was measured using the CellTiterGlo assay according to the manufacturer’s instructions, and luminescence was recorded with a microplate reader. Cell viability was calculated as: cell viability (%) = RLU_sample_ / RLU_cell only_ × 100%.

### Virus challenge with MERS-CoV in human *ex vivo* lung tissues

Human lung tissues for *ex vivo* studies were retrieved from patients undergoing surgical operations at the Queen Mary Hospital, Hong Kong. All donors gave written consent as approved by the Institutional Review Board of the University of Hong Kong/Hospital Authority Hong Kong West Cluster. Normal nonmalignant lung tissue fragments in excess for clinical diagnosis were used. The freshly obtained lung tissues were processed into small rectangular pieces and were rinsed with the primary tissue culture medium, which contained the advanced DMEM/F12 medium supplemented with 2 mM HEPES (Gibco), 1X GlutaMAX (Gibco), 100 U/mL penicillin, 100 μg/mL streptomycin, 20 μg/mL vancomysin, 20 μg/mL ciprofloxacin, 50 μg/mL amikacin, and 50 μg/mL nystatin as previously described(*71, 72*). The specimens were infected with MERS-CoV at a titre of 3×10^6^ PFU in 1 mL culture medium. After 2 h, the inoculum was removed and the specimens were washed thoroughly with the primary tissue culture medium. The infected tissues were then incubated with primary tissue culture medium. Tissues were collected at 24 hpi for TCID_50_ or fixed with 4% paraformaldehyde for immunofluorescence staining.

### Cell lines

HT29 cells obtained from ATCC were maintained in McCoy’s 5A medium (GibcoTM, 16600082) supplemented with 10% heat-inactivated fetal bovine serum (FBS), 100 U/mL penicillin, and 100 μg/mL streptomycin. HT29-ACE2 and HT29 knockout cells were maintained in McCoy’s 5A medium supplemented with 10% FBS, 100 U/mL penicillin, 100 μg/mL streptomycin, and 5 μg/mL puromycin. A549, Huh7, 293T, VeroE6, Caco2, MRC5 and LLK-MK2 cells obtained from ATCC were maintained in Dulbecco’s Modified Eagle Medium (DMEM) (GibcoTM, 11965092) supplemented with 10% FBS, 100 U/mL penicillin, and 100 μg/mL streptomycin. VeroE6-TMPRSS2 cells were obtained from the Japanese Collection of Research Bioresources (JCRB) Cell Bank and cultured in DMEM with 10% FBS, 100 U/mL penicillin, 100 μg/mL streptomycin, and 1mM (1%) sodium pyruvate. BEAS2B and Calu3 cells obtained from ATCC were maintained in DMEM/F12 (GibcoTM, 12634010) supplemented with 10% FBS, 100 U/mL penicillin and 100 μg/mL streptomycin. Human primary monocytes and T cells were obtained from human peripheral blood mononuclear cells (PBMCs) taken from healthy donors, collected from Hong Kong Red Cross Blood Transfusion Service according to a protocol approved by the Institutional Review Board of The University of Hong Kong. MDMs were differentiated from monocytes in Roswell Park Memorial Institute (RPMI) 1640 Medium (GibcoTM, 12633012) supplemented with 10% FBS, 100 U/mL penicillin, 100 μg/mL streptomycin, 2 mM glutamine, 1% sodium pyruvate, 1% non-essential amino acids, and 10 ng/mL recombinant human granulocyte macrophage colony-stimulating factor (GM-CSF) (R&D Systems).

### Chemical modulators

zVAD-FMK (HY-16658B), Necrostatin-1 (Nec-1) (HY-15760), GSK872 (HY-101872), GSK840 (HY-104021), Necrosulfonamide (NSA) (HY-100573), BV6 (HY-16701) and HS-15 (HY-Y1893) were obtained from MedChemExpress. Human TNF-α was obtained from R&D Systems (210-TA). UH15-38 was provided by S. Balachandran.

### Antibodies for Western blots

MERS-CoV N was detected by an in-house guinea pig immune serum (1:100,000). Anti-ZBP1 (A04739-2, 1:2,000) was from Boster. Anti-RIPK1 (ab300617, 1:2,000) and anti-caspase-3 (ab32351, 1:5,000) were from Abcam. Anti-pRIPK1 (28252-1-AP, 1:2,000), anti-RIPK3 (17563-1-AP, 1:10,000) and anti-MLKL (66675-1-Ig, 1:10,000) were from Proteintech. Anti-pRIPK3 (93654S, 1:2,000) was from CST. Anti-pMLKL (14251, 1:2,000) was from SAB. Anti-flag (M185-3L, 1:10,000) and anti-myc (M192-3, 1:10,000) were from MBL. Anti-β-actin (AC028, 1:10,000) was from Abclonal. Secondary antibodies, including goat anti-mouse horseradish peroxidase (HRP) (62-6520, 1:5,000), goat anti-rabbit HRP (65-6120, 1:5,000), and goat anti-guinea pig HRP (A18769, 1:10,000) were from Thermo Fisher Scientific. HRP conjugated affiniPure goat anti-mouse IgG (H+L) (BA1050, 1:8,000) and HRP conjugated affiniPure goat anti-rabbit IgG (H+L) (BA1055, 1:8,000) were from Boster.

### Antibodies for immunofluorescence staining

MERS-CoV N was detected by an in-house guinea pig immune serum (1:2,000). SARS-CoV-1 and SARS-CoV-2 N was detected by an in-house rabbit immune serum (1:1,000) or mouse anti-SARS-CoV-2 N (ab281300, 1:1,000) from Abcam. HCoV-NL63 N was detected by an in-house rabbit immune serum (1:1,000). Anti-HCoV-OC43-N (40643-T62, 1:1,000 and anti-HCoV-229E-N (40640-T6, 1:1,000) were from Sino Biological. Anti-ZBP1 (A04739-2, 1:200) was from Boster. Anti-Z-RNA (Z22, 1:200) was from Absolute. Anti-MLKL (phospho S358) antibody (ab187091, 1:1,000) was from Abcam. Anti-flag (AE092, 1:500) was from Abclonal. Anti-myc (M192-3, 1:500) was from MBL. Secondary antibodies, including Alexa Fluor 488 goat anti-guinea pig IgG (H+L) (A11073, 1:1,000), Alexa Fluo 568 goat anti-mouse IgG (H+L) (A11004, 1:1,000), Alexa Fluor 594 goat anti-mouse IgG (H+L) (A11005, 1:1,000), Alexa Fluor 647 goat anti-rabbit IgG (H+L) (A21245, 1:1,000), Alexa Fluor 488 goat anti-mouse IgG (H+L) (A11001, 1:1,000), Alexa Fluor 594 goat anti-rabbit IgG (H+L) (A11012, 1:1,000), Alexa Fluor 488 donkey anti-rabbit IgG (H+L) (A21206, 1:1,000), Alexa Fluor 594 donkey anti-rabbit IgG (H+L) (A21207, 1:1,000) were from Thermo Fisher Scientific.

### Antibodies for flow cytometry

MERS-CoV N was detected by an in-house guinea pig immune serum (1:100). HCoV-NL63 N was detected by an in-house rabbit immune serum (1:100). Antibody against Z-RNA (Ab00783-3.0, 1:100) was from Absolute antibody. pMLKL (Phospho-Ser345) antibody (14251, 1:100) was from SAB.

### HT29 knockout cell line construction

HT29 knockout cells were generated using the lentiCRISPRv2 vector (Addgene, 52961) to target the gene of interest. To clone the guide sequence into lentiCRISPRv2, two complementary oligonucleotides corresponding to the target sequence were synthesized. Each oligonucleotide pair was phosphorylated and annealed using T4 polynucleotide kinase (T4 PNK, NEB, M0201S). The annealed oligonucleotides were then ligated into the BsmBI-digested lentiCRISPRv2 plasmid. Lentivirus was produced by co-transfecting HEK-293T cells with the sgRNA-containing lentiCRISPRv2 plasmid, psPAX2 (Addgene, 12260) and pVSVg (Addgene, 8454). Viral supernatants were collected 48 h after transfection. HT29 cells were transduced with lentiviral supernatants and selected with puromycin (10 μg/mL) for 7 days. Surviving cells were collected, dissociated into single-cell suspensions and single cell sorting by flow cytometry (BD FACSAria Fusion/SORP). Monoclonal cell lines were expanded, and whole-cell lysates were prepared in RIPA buffer. Protein expression in the knockout cells were confirmed by Western blots.

### Cell death modulators and virus challenge

HT29 or HT29 knockout cells were mock-infected or infected with MERS-CoV at 10 MOI. The infected cells were treated with 30 ng/mL human TNF-α, 1 μM BV6, 25 μM zVAD-FMK, 25 μM Necrostatin-1, 5 μM GSK872, 3 μM GSK840, 5 μM NSA, DMSO, or a combination of the above-mentioned cell death modulators. At 24 hpi, total RNA from cell lysate was extracted for RT-qPCR analysis of MERS-CoV N expression. The cell viability of treated and virus infected cells were quantified using CellTiterGlo assays (Promega, G7570) and signals were recorded by PerkinElmer EnSight Multimode Plate Reader or GloMax Explorer Multimode Microplate Reader. Cellular morphology was documented by microscopy. To quantify virus induced-pMLKL expression, HT29-ACE2 cells were challenged with MERS-CoV (1 MOI), SARS-CoV-1 (1 or 5 MOI), or SARS-CoV-2 (1 or 5 MOI). At the indicated time points, pMLKL was detected by enzyme-linked immunosorbent assay (ELISA) (Abcam, ab279863) using the GloMax Explorer Multimode Microplate Reader.

### MERS-CoV challenge with cell death modulator treatment in human intestinal organoids

Human intestinal organoids were established using biopsied human intestinal tissues from patients who underwent surgical operations at the Queen Mary Hospital, Hong Kong. All donors had written consent as approved by the Institutional Review Board of the University of Hong Kong/Hospital Authority Hong Kong West Cluster. Human intestinal organoids were established and maintained in expansion medium, and differentiation was induced by incubating with differentiation medium for 5 days as previously described(*73–75*). Differentiated intestinal organoids were sheared mechanically and inoculated with MERS-CoV at 0.1 MOI for 2 h. After the inoculum was removed, the intestinal organoids were rinsed with basal medium (Advanced DMEM/F-12 (Gibco), 1% HEPES, 1% GlutaMAX, and 1% Penicillin/Streptomycin) and embedded in Matrigel and maintained in differentiation medium containing 30 ng/mL human TNF-α, 1 μM BV6, 25 μM zVAD-FMK, 5 μM NSA, DMSO, or a combination of the above-mentioned cell death modulators. At 24 hpi, cell viability was quantified with CellTiterGlo assays and cellular morphology was documented by microscopy.

### RNA extraction and one-step reverse transcription-quantitative polymerase chain reaction (RT-qPCR)

RNA of homogenized animal tissues or cell lysates was extracted by using E.Z.N.A. Total RNA Kit I (Omega, R6834-03). Quantitative reverse transcription polymerase chain reaction (RT-qPCR) was conducted with UniPeak U+ One Step RT-qPCR SYBR Green Kit (Vazyme, Q226-01). Quantitative reverse transcription PCR was performed on LightCycler 480 Real-Time PCR System (Roche, Basel, Switzerland) to detect the expression of viral gene copies or immune response related genes.

### Infectious titer quantification

Infectious titer of infected cells and infected animal lung tissues were determined by TCID50. In brief, the collected supernatant samples were serially diluted and inoculated to VeroE6 or VeroE6-TMPRSS2 cells in 96-well plates for 2 h at 37 °C. After inoculation, the residual viral particles were removed, and cells were washed three times with serum-free DMEM. At four days post-infection, cytopathic effects (CPE) were evaluated using the Muench and Reed method.

### Western blots

Cells were lysed with RIPA buffer (Byotime, P0013C) with protease inhibitor (MCE, HY-K0010), followed by boiling for 10 min in SDS-PAGE loading buffer (EpiZyme, LT101). Proteins were separated by 8-12% SDS–PAGE gels and transferred to PVDF membranes (Vazyme, E801-01; Bio-Rad, 1620177). Membranes were blocked with PBST (MCE, HY-K1022) containing 5% (w/v) non-fat dry milk at room temperature for 1 to 4 h, and incubated with specific primary antibodies overnight at 4 °C, followed by HRP-conjugated secondary antibodies for 45 min to 1 h at room temperature. The signal was visualized using HRP substrate (Merck Millipore, WBLUF0500) and detected with an Alliance Imager apparatus (Uvitec).

### Flow cytometry

Cells were washed with PBS and stained with Zombie Aqua Fixable Viability Kit (BioLegend 423101, 1:500) according to the manufacturer’s instructions. After viability stain, cells were fixed with 4% paraformaldehyde, permeabilized with 90% methanol, and stained intracellularly with antibodies against viral N, cleaved caspase-3, pMLKL, or Z-RNA as indicated for each experiment. All samples subjected to Z-RNA staining were treated with DNaseI (TakaRa, 2270A, 25 U/mL) before anti-Z-RNA labelling, to reduce DNA-dependent background signals. Flow cytometry was performed using a BD FACSymphony A1 flow cytometer and data were analyzed using FlowJo version 10.10.0 (BD Biosciences). Single cells were gated by FSC-A/FSC-H, followed by exclusion of Zombie-positive dead cells. Positive gates were set using mock-infected, unstained and single-stained controls. The gating strategies were demonstrated in **Fig. S9**.

### Lung tissue histological and immunofluorescence staining

Infected lung tissues were fixed in 4% paraformaldehyde. A Leica TP1020 processor was used to conduct ethanol-washing, and paraffin-embedding. To conduct histological and immunofluorescence staining, tissues was sectioned with a microtome-cut (Thermo Fisher) and oven-dried (37 °C overnight). For immunofluorescence staining, slides underwent antigen unmasking (Vector Labs, H-3300), Sudan black B quenching, and blocking before primary antibody incubation. Anti-MERS-CoV N, anti-SARS-CoV-1 N, anti-SARS-CoV-2 N, and anti-pMLKL antibodies were applied to visualize viral or pMLKL protein. Alexa Fluor secondary antibodies were used followed by DAPI mounting (Vector H-1200). Images were captured with a Carl Zeiss LSM800 confocal microscope using ZEISS (ZEN 2) software. For H&E staining, the tissue sections were stained with Gill’s haematoxylin (Vector Laboratories, H-3401-500) and eosin-Y. All images were captured with an Olympus MDX10 microscope (Olympus Life Science) or an Olympus BX53 fluorescence microscope (Olympus Life Science). H&E-stained lung tissue sections were blinded for the identities of experimental groups and examined by a trained histopathologist. Lung pathology was graded on a scale of 0 (normal) to 4 (most severe) adapted from a grading system we previously described(*12, 13*). In brief, lung injury was assessed using a semi-quantitative scoring system with severity graded according to four key pathological changes including inflammatory cell infiltration, alveolar space exudation, alveolar wall collapse, and epithelium damage. The scoring criteria were defined as: 0, normal lung parenchyma with no significant pathological changes; 1, vascular congestion with perivascular or peribronchial inflammatory cell infiltration; 2, diffuse alveolar wall congestion and inflammatory infiltration, along with early alveolitis, extending beyond the changes seen in score 1; 3, intra-alveolar inflammatory exudation, cellular infiltration, and focal alveolar wall collapse; 4, diffuse alveolitis with widespread destruction of alveolar architecture and marked epithelial injury.

### Immunofluorescence staining of cell line samples

Immunofluorescence staining was performed as we previously described(*76, 77*). In brief, cell monolayers seeded in chamber slide (SPL, 30108) were immobilized with 4% paraformaldehyde at room temperature for 20 min and washed 3 times with PBS. Next, the cells were permeabilized with 0.1% Triton-X 100 in PBS for 5 to 10 minutes at room temperature and blocked with blocking buffer (10% goat serum diluted with PBS) for 1.5 h at room temperature. Cells were then incubated with primary antibodies for 1.5 h and secondary antibodies for 45 min at room temperature, followed by staining for DAPI (Beyotime, C1006-50ml) for 10 mins at room temperature. When required, DNase I (Takara, 2270A, 5 U/mL or 25 U/mL), RNase A (Beyotime, ST576, 1 mg/mL) and RNase III (Beyotime, R7086M, 50 U/mL) were applied for 1 h at 37°C before primary antibodies treatment. Chamber slides were covered with Prolong Diamond Antifade mountant solution (Invitrogen, P36970) to prevent quenching, and imaged with an Olympus BX53 fluorescence microscope (Olympus Life Science), Carl Zeiss LSM800 or LSM900 confocal microscope.

### Biological sample processing, single-cell library construction, and sequencing

A total of 24 bronchoalveolar lavage fluid specimens were collected from 24 participants, comprising 16 patients infected with the SARS-CoV-2 Omicron variant and 8 control subjects. Among the 16 patients, 9 survived and 7 died during follow-up. Bronchoalveolar lavage fluid (BALF) specimens from patients were obtained during bronchoalveolar lavage performed for clinically indicated diagnostic or therapeutic purposes. The study was approved by the Ethics Committee of China-Japan Friendship Hospital (2022-KY-052-1), and written informed consent was obtained from all participants or their legal representatives. After collection, BALF samples were gently filtered through a 70 μm strainer to eliminate mucus, debris, and cell aggregates. The filtrates were then centrifuged at 300 ×g for 5 min at 4 °C, and the resulting cell pellets were retained for subsequent processing. Following erythrocyte depletion with red blood cell lysis buffer, cell concentration and viability were determined by AO/PI staining using a Countstar Rigel S2 fluorescence cell analyzer. For samples with suboptimal viability, dead cells were depleted using a magnetic bead-based dead cell removal procedure. The purified cells were washed twice with RPMI 1640 medium and resuspended in RPMI 1640 containing 2% FBS. The final cell density was adjusted to 1×10^6^ cells/mL before single-cell capture. Cell barcoding and 3’ single-cell transcriptome library preparation were performed using the SeekOne Digital Droplet Single Cell 3’ system, including the corresponding library preparation reagents, barcoded hydrogel beads, and SeekOne S3 chip, in accordance with the manufacturer’s protocol. The completed libraries were subjected to paired-end sequencing on an Illumina NovaSeq 6000 platform with a read length of 150 bp.

### Processing and quality control of single-cell RNA-seq data

Sequencing reads were converted into gene-by-cell count matrices using SeekSoul Tools after alignment to the human GRCh38 reference genome. Putative doublets were evaluated separately for each sample with Scrublet. Downstream quality control was conducted in Scanpy v1.9.3. Cells were retained only when at least 200 genes were detected, whereas genes expressed in fewer than three cells were discarded. Cells with mitochondrial transcripts representing more than 10% of total UMI counts were also excluded. The filtered matrices were normalized, followed by log1p transformation. Highly variable genes were selected for downstream dimensionality reduction, and the effect of total UMI counts was regressed out before principal component analysis (PCA). After PCA, all samples were projected into a shared analytical space and subjected to unsupervised clustering. Harmony was then used with default parameters to reduce inter-sample batch effects before subsequent visualization and downstream analyses.

### Sequence data collection for GC content ratio and Z-RNA site calculation

Viral genomic sequences representing major human-pathogenic coronaviruses and bat coronaviruses were retrieved from GenBank. The initial dataset comprised seasonal human coronaviruses (1,028 HCoV-HKU1; 1,563 HCoV-NL63; 2,413 HCoV-OC43; 1,401 HCoV-229E), epidemic and pandemic coronaviruses (765 MERS-CoV; 140 SARS-CoV-1; 2,998 SARS-CoV-2), and 185 distinct bat coronaviruses. To ensure sequence quality, genomes containing >20% ambiguous nucleotides (Ns) were excluded from the analysis. Following this filtering step, the final dataset consisted of 361 HCoV-HKU1, 402 HCoV-NL63, 807 HCoV-OC43, 551 HCoV-229E, 122 SARS-CoV, 339 MERS-CoV, 2,885 SARS-CoV-2, and 111 Bat CoV sequences. For the longitudinal analysis of SARS-CoV-2, we retained 2,998 GenBank sequences sampled between the pandemic’s onset and May 1, 2026. Sequence selection was guided by the representative Nextstrain global phylogenetic tree (https://nextstrain.org/ncov/open/global/all-time@2026-05-01) utilizing a global subsampling strategy. After removing outliers, to map the virus’s evolutionary trajectory, 2,885 sequences were categorized by clade to trace the shift from historical variants (Alpha, Beta, Gamma, and Delta) to contemporary Omicron sublineages (e.g., BA.4, BA.5, BQ.1, XBB.1.5, EG.5.1, JN.1, KP.3, XEC, XFG, and BA.3.2). The final dataset comprises 52 distinct clades (ranging from ancestral 19A and 19B to the recent 25G, XFC) containing 1 to 411 sequences per clade, and encompasses 677 unique Pango lineages with 1 to 167 sequences per lineage.

### GC content and Z-RNA site calculation

For each viral genome, the overall GC content ratio (GC%) was calculated by dividing the total count of guanine (G) and cytosine (C) nucleotides by the valid sequence length (defined as the sum of standard A, T, G, and C bases). The propensity for sequences to adopt the Z-conformation (Z-score) was evaluated using the **zhunt3** algorithm(*43, 44*). The algorithm was executed across the full viral genomes to identify potential Z-forming sequences ranging from 6 to 24 nucleotides in length. To isolate high-confidence Z-forming sites from the algorithm’s output, a strict Z-score threshold of >1.0 was applied. Only loci meeting or exceeding this propensity score were counted as favorable for Z-conformation. To account for variations in genome size across different viral strains and classes, the raw count of predicted Z-forming sites was normalized per 100 base pairs (bp). Ambiguous nucleotides (Ns) were explicitly excluded from the total length during this calculation. This normalization provided a standardized metric, allowing for accurate comparative longitudinal and cross-lineage analyses. To assess the evolutionary trajectory of Z-RNA forming potential in SARS-CoV-2, a temporal trend analysis was conducted. Linear regression was applied to the Z-RNA propensity values plotted against sequence collection dates spanning 2020 to 2026. The strength and statistical significance of these temporal correlations were evaluated using the coefficient of determination (R^2^) and *P*-values.

### Fluorescence intensity analysis

Fluorescence intensity of CoV N, pMLKL, and Z-RNA, were determined in human coronavirus-infected tissues by ImageJ. Measurements were conducted in no less than three randomly selected microscopic fields and the n numbers were indicated in the corresponding figure legends. N fluorescence intensity was taken as the internal reference to calculate the relative ratio of pMLKL to N, or Z-RNA to N.

### Statistical analysis

Data in figures are mean ± standard error of the mean (SEM). Statistical comparison between different groups was performed by one-way ANOVA, two-way ANOVA, Student’s t-test or Log-rank (Mantel-Cox) test using GraphPad Prism 10. Differences were considered statistically significant at *P* < 0.05.

## Supplementary Materials

Figs. S1 to S9 for multiple supplementary figures.

## Acknowledgements

We thank the Imaging and Flow Cytometry Core (CPOS) and Centre for Comparative Medicine Research at The University of Hong Kong for their supports on data collection.

## Funding

Prevention and Control of Emerging and Major Infectious Diseases-National Science and Technology Major Project (2026ZD01911200)

National Natural Science Foundation of China (82502687, 92369201)

Health and Medical Research Fund (CID-HKU1-5, 25240662, 23220522, 20190652), the Food and Health Bureau, The Government of the Hong Kong Special Administrative Region

General Research Fund (17118621, 17119122), Collaborative Research Fund (C7103-22G and C7060-21G), and Theme-Based Research Scheme (T11-709/21-N), the Research Grants Council of the Hong Kong Special Administrative Region

The InnoHK initiative of the Innovation and Technology Commission of the Hong Kong Special Administrative Region Government

Seed Fund for Collaborative Research

National Program on Key Research Project of China (2020YFA0707500, 2020YFA0707504)

Guangdong Provincial National Science Foundation, China (2023A1515012325, 2024A1515010848)

The Major Science and Technology Program of Hainan Province (ZDKJ202003)

The research project of Hainan Academician Innovation Platform (YSPTZX202004); the Hainan Talent Development Project (SRC200003)

The High Level-Hospital Program, Health Commission of Guangdong Province, China

Emergency COVID-19 grant (2021YFC0866100) from Major Projects on Public Security under the National Key Research and Development Program of China

National Natural Science Foundation of China Young Student Basic Research Program (825B2063 to Y.L.)

National Natural Science Foundation of China Young Scientist Fund Type (A) (82525037 to J.F.-W.C.)

Research Fellow Scheme (HKU RFS2425-7S06 to J.F.-W.C), Research Grants Council of the Hong Kong Special Administrative Region, China

Major Projects of Guangzhou National Laboratory (GZNL2025C01040, GZNL2024A01017 and GZNL2023A01009)

National Science Foundation of China (32350710189 to FKMC) Key R&D Program of Zhejiang Province, China (to FKMC)

Support from Dr. Gallant Ho on the Gallant Ho Outstanding Young Professorship.

Donations of the Shaw Foundation Hong Kong, Richard Yu and Carol Yu, May Tam Mak Mei Yin, Michael Seak-Kan Tong, Yip Mow Lum, the Providence Foundation Limited (in memory of the late Lui Hac Minh), Lee Wan Keung Charity Foundation Limited, Hui Ming, Hui Hoy and Chow Sin Lan Charity Fund Limited, Hong Kong Sanatorium & Hospital, The Chen Wai Wai Vivien Foundation Limited, Chan Yin Chuen Memorial Charitable Foundation, Marina Man-Wai Lee, the Hong Kong Hainan Commercial Association South China Microbiology Research Fund, the Jessie & George Ho Charitable Foundation, Perfect Shape Medical Limited, Kai Chong Tong, Foo Oi Foundation Limited, Tse Kam Ming Laurence, Betty Hing-Chu Lee, Ping Cham So, and Lo Ying Shek Chi Wai Foundation.

All donors and funding sources had no role in the study design, data collection, analysis, interpretation, or writing of the manuscript.

## Author contributions

Conceptualization: HC, KYY

Methodology: HC, YHou, JS, BS, MZ, HZ, YFH, HL, ZS, YHe, ZC, CD, HL, JL, AG

Investigation: YHou, JS, BS, MZ, HZ, YFH, HL, ZS, YHe, ZC, CD, HL, JL, AG, CL, LW, CY, YC, YW, TZ, YL, BH, TTTY, XZ, JC, HS, CL, XZ, ML, XM, SY, JZ, BZZ, KFLL, ZW, KHK, JDH, DYJ

Visualization: HC, YHou

Funding acquisition: HC, KYY, SB, FKMC, BC

Project administration: HC, KYY, SB, FKMC, BC, YHou Supervision: HC, KYY, SB, FKMC, BC

Writing – original draft: HC, YHou

Writing – review & editing: HC, KYY, SB, FKMC, BC, YHou, JS, BS, MZ, YFH, HL, HZ

## Data and materials availability

All data are available in the main text or the supplementary materials

## Supplementary Materials

**Fig. S1.**
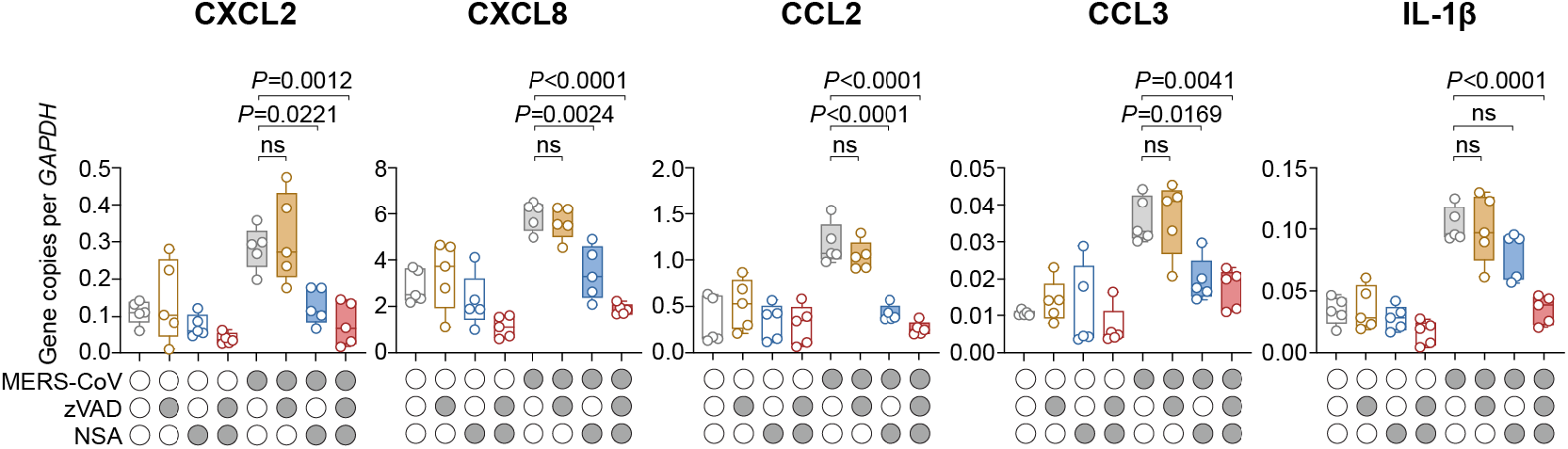
Necroptosis inhibition modulates MERS-CoV-induced inflammation. MERS-CoV-or mock-infected human *ex vivo* lung tissues were treated with 50 μM zVAD, 5 μM NSA, or a combination of both. Tissue lysate samples were harvested at 24 hpi to determine the expression of pro-inflammatory markers (IL-1β) and inflammatory cell-recruiting chemoattractant (CXCL2, CXCL8, CCL2, CCL3) (n=5). Data represented median and interquartile range from the indicated number of biological repeats. Statistical significance between groups was determined with one way-ANOVA with Tukey’s multiple comparisons tests.

**Fig. S2.**
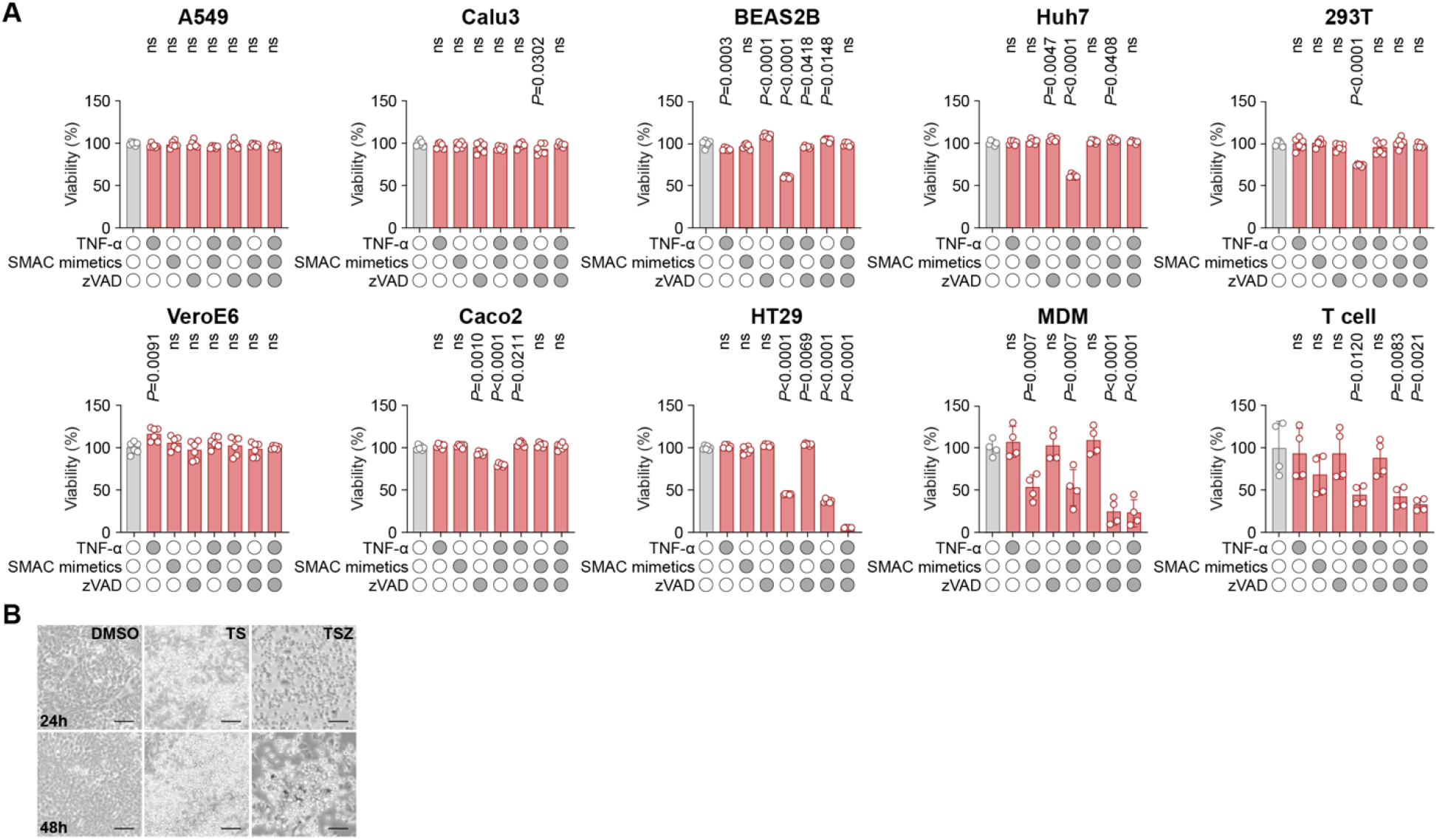
Necroptosis sensitivity in conventional cell lines used in coronavirus research. (**A**) Commonly used cell types for coronavirus studies were treated with the indicated combinations of 30 ng/mL TNF-α (T), 1 μM smac mimetic (BV6) (S), and 25 μM zVAD (Z). Cell viability was quantified with CellTiterGlo assays at 24 hours post treatment. (**B**) Representative images of HT29 cells treated with TS or TSZ at 24 and 48 hours post treatment. Data represented mean and standard error of the mean from the indicated number of biological repeats. Statistical significance between groups was determined with one way-ANOVA with Dunnett’s multiple comparisons tests.

**Fig. S3.**
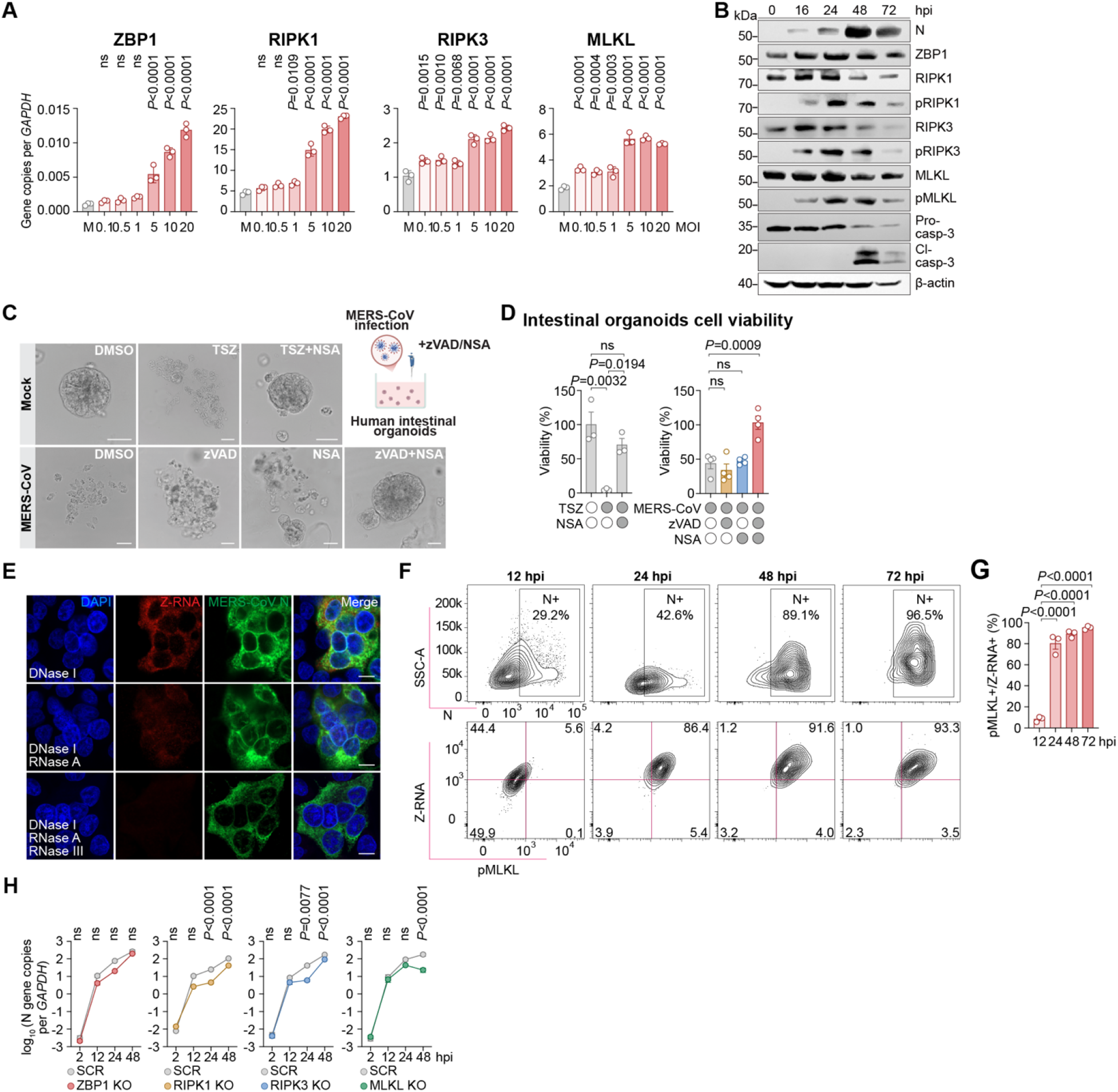
Mechanistic investigation on coronavirus-induced necroptosis. (**A**) HT29 cells were infected with MERS-CoV at the indicated MOI for 24 h. The expression of ZBP1, RIPK1, RIPK3, and MLKL was quantified with RT-qPCR. Gene expression was normalized with the expression of GAPDH (n=3). (**B**) HT29 cells were infected with MERS-CoV at 1 MOI. Cell lysates were harvested at the indicated time points post infection for Western blot analysis. (**C-D**) Human intestinal organoids were infected with MERS-CoV at 0.1 MOI and treated with 50 μM zVAD, 5 μM NSA, or a combination of both (n=4). Mock-infected intestinal organoids treated with TSZ or TSZ/NSA were included as controls (n=3). Representative images were recorded at 24 hpi (C). Cell viability was quantified with CellTiterGlo assays at 24 hpi (D). (**E**) HT29 cells were infected with MERS-CoV at 1 MOI. At 20 hpi, infected cells were fixed and treated with DNase I (25 U/mL), RNase A (1 mg/mL), RNase III (50 U/mL), in single or combined treatments for 1 h at 37°C. Z-RNA and MERS-CoV N were immunolabelled and revealed with confocal microscopy. (**F-G**) HT29 cells were infected with MERS-CoV at 4 MOI and fixed at the indicated time points. Z-RNA, pMLKL, and MERS-CoV N were immunolabelled and analyzed with flow cytometry (n=3). (**H**) ZBP1, RIPK1, RIPK3, or MLKL KO HT29 cells were infected with MERS-CoV at 10 MOI and harvested at the indicated time points. MERS-CoV N gene expression was quantified with RT-qPCR after normalizing with GAPDH expression. MERS-CoV N gene expression in scrambled KO cells was determined and used as the control (n=4). Data represented mean and standard error of the mean from the indicated number of biological repeats. Statistical significance between groups was determined with one way-ANOVA with Dunnett’s multiple comparisons test (A), one way-ANOVA with Tukey’s multiple comparisons test (D,G) or two way-ANOVA with Šídák’s multiple comparisons test (H). Bars in (C,E) represented 50 μm and 10 μm, respectively.

**Fig. S4.**
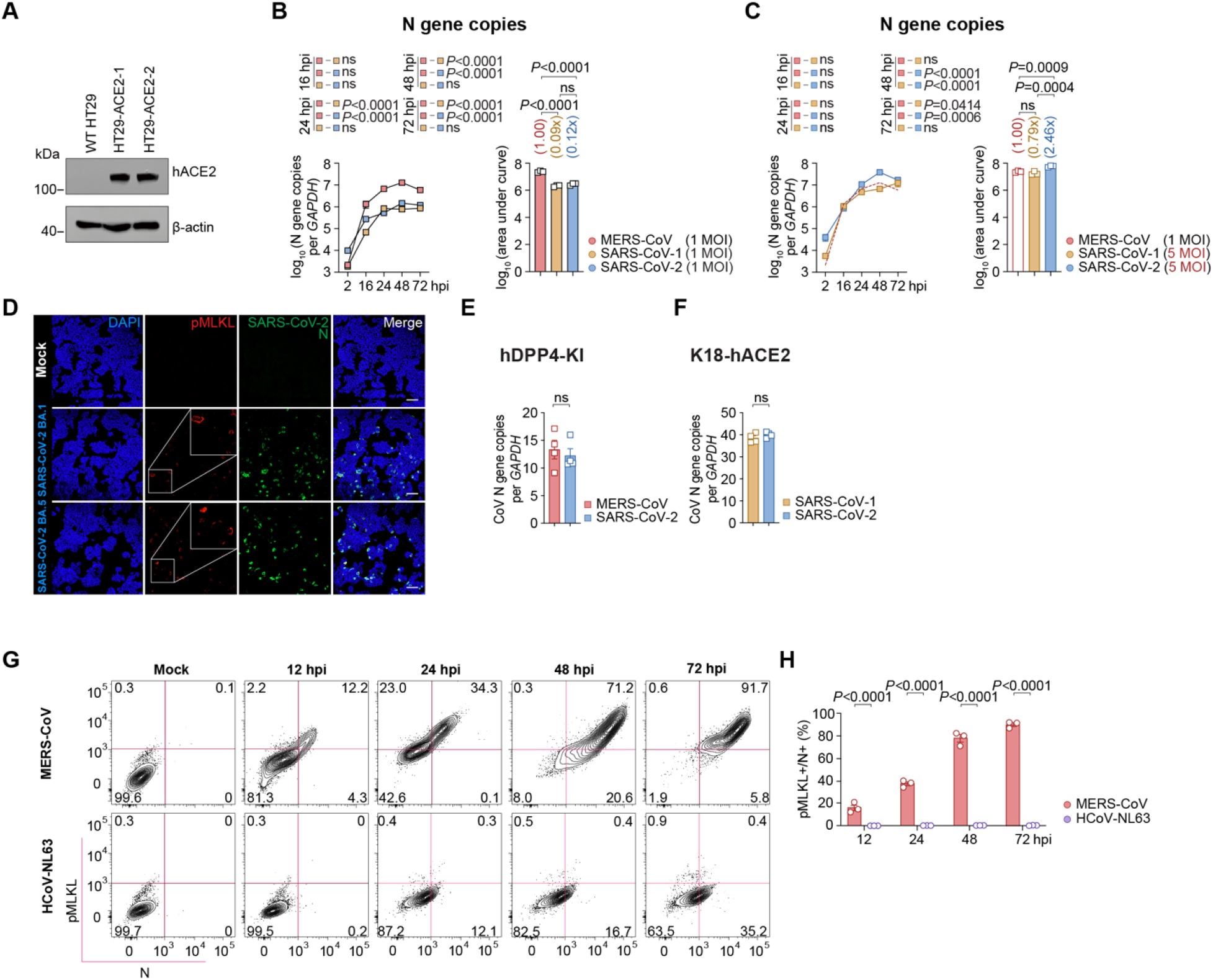
Replication of MERS-CoV, SARS-CoV-1, and SARS-CoV-2 in vitro and in vivo. (**A**) Western blot for human ACE2 expression in HT29 and HT29-ACE2 cells. (**B-C**) HT29-ACE2 cells were infected by MERS-CoV, SARS-CoV-1, or SARS-CoV-2 at the indicated MOI. Cell lysates were harvested from 2 to 72 hpi. Virus replication was quantified with RT-qPCR. (**D**) HT29-ACE2 cells were infected with SARS-CoV-2 Omicron BA.1 or BA.5 at 10 MOI. At 24 hpi, infected cells were fix for immunofluorescence staining and confocal microscopy to visualize pMLKL and N protein expression. (**E-F**) N gene expression of coronaviruses in lungs of hDPP4-KI and K18-hACE2 mice were assessed with RT-qPCR at 3 dpi (n=4). (**G-H**) HT29-ACE2 cells were infected with 1 MOI MERS-CoV or 10 MOI HCoV-NL63. The infected cells were fixed at the indicated time points and immunolabelled for N and pMLKL expression for flow cytometry analysis (G). Cells double positive for N and pMLKL were summarized (n=3) (H). Data represented mean and standard error of the mean from the indicated number of biological repeats. Statistical significance between groups was determined with one way-ANOVA with Tukey’s multiple comparisons tests and two-way ANOVA with Tukey’s multiple comparisons tests (B,C) or two-tailed Student’s t-tests (E,F). Bars in (D) represent 100 μm.

**Fig. S5.**
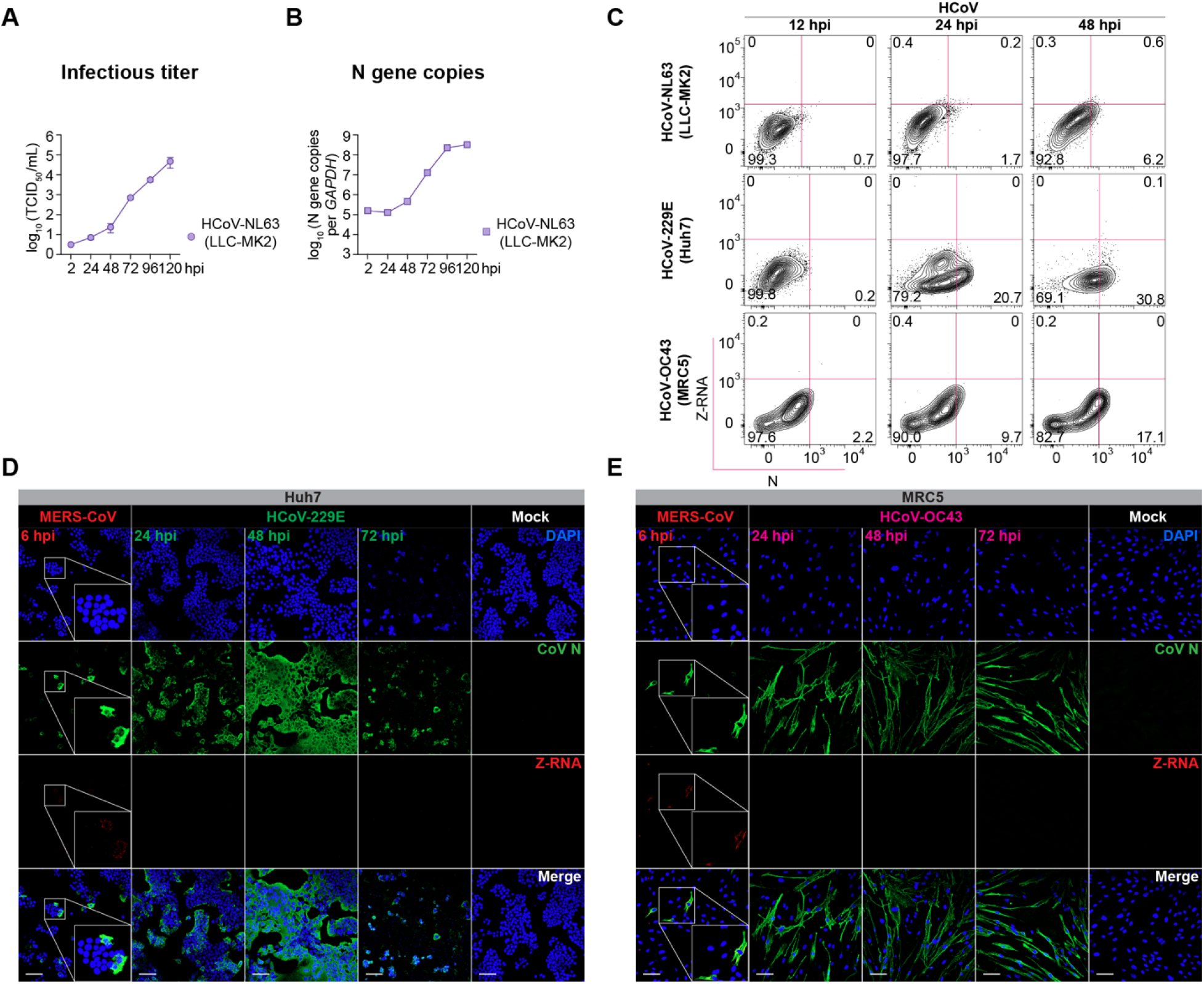
Differential Z-RNA induction between highly pathogenic and seasonal coronaviruses. (**A-B**) LLC-MK2 cells were infected with HCoV-NL63 at 1 MOI. Infected cells were harvested at the indicated time points and virus replication was determined with TCID_50_ assays (A) or RT-qPCR (B). (**C**) Seasonal coronavirus-infected cells were fixed at the indicated time points for detection of Z-RNA and N protein with flow cytometry. (**D**) Huh7 cells were infected with HCoV-229E at 5 MOI and fixed at 24, 48, and 72 hpi. As a positive control, Huh7 cells were infected with MERS-CoV at 1 MOI and fixed at 6 hpi. Mock-infected cells were included as negative controls. (**E**) MRC5 cells were infected with HCoV-OC43 at 5 MOI and fixed at 24, 48, and 72 hpi. As a positive control, MRC5 cells were infected with MERS-CoV at 1 MOI and fixed at 6 hpi. Mock-infected cells were included as negative controls. Z-RNA and coronavirus N protein expression were revealed with immunofluorescence staining and confocal microscopy. Cell nuclei were revealed with DAPI staining. Data represented mean and standard error of the mean from the indicated number of biological repeats. Bars in (D,E) represents 100 μm.

**Fig. S6.**
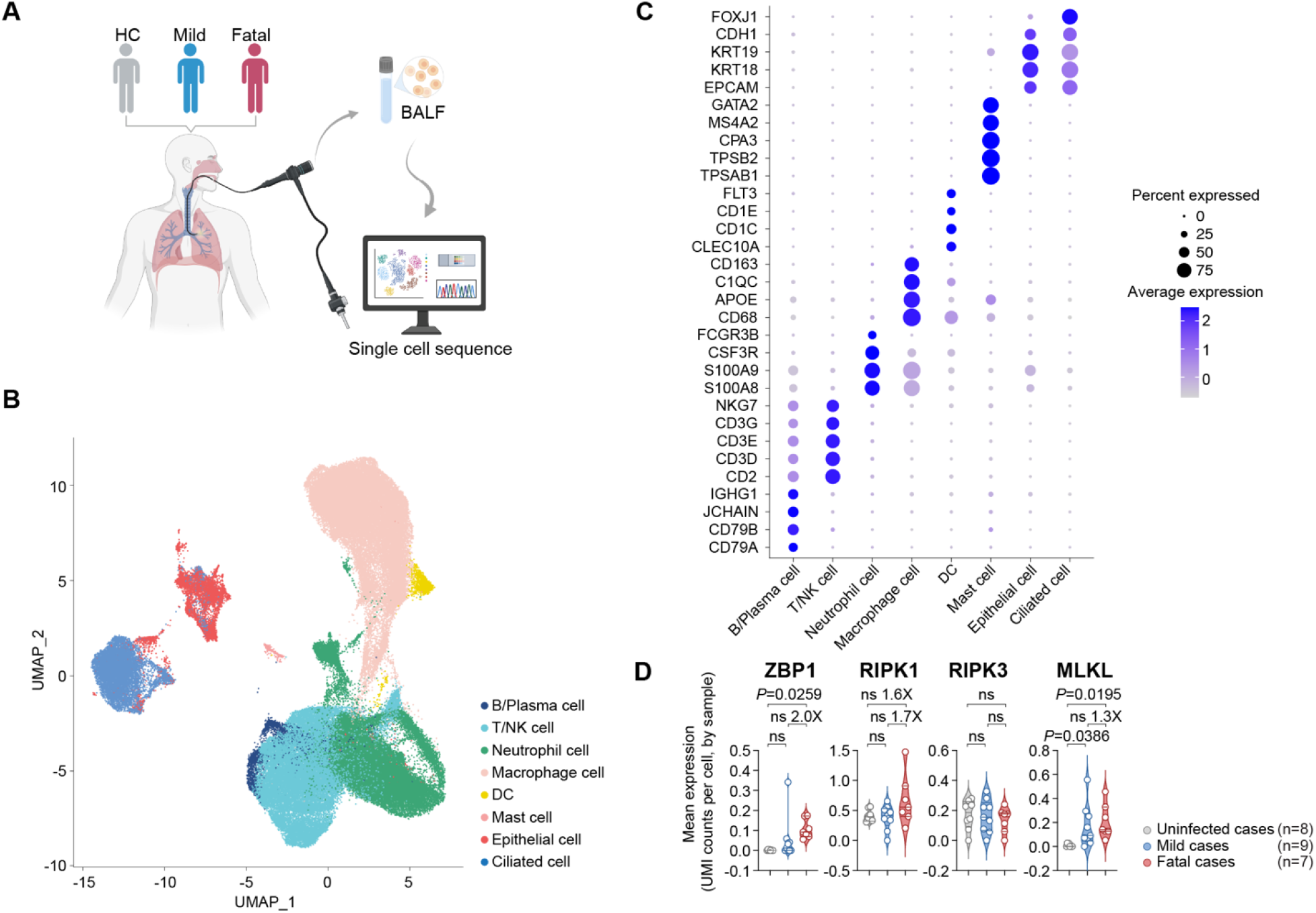
ZBP1 is significantly upregulated in COVID-19 patients with fatal outcomes. (**A**) Schematic of sample collection. (**B**) UMAP visualization of all analysed cells colored by annotated cell type. Major cell populations included B cells, plasma cells, T/NK cells, neutrophils, macrophage cells, DC, epithelial cells, and ciliated cells. Each dot represents a single cell, and colors indicate cell-type annotation. (**C**) Dot plot that shows the expression of representative canonical gene markers used for cell-type annotation. B cells were characterized by CD79A and CD79B; plasma cells by JCHAIN, IGKC, and IGHG1; T/NK cells by CD2, CD3D, CD3E, CD3G, and NKG7; neutrophils by S100A8, S100A9, FCGR3B and CSF3R; macrophages by CD68, APOE, C1QC, and CD163; dendritic cells by CLEC10A, CD1C, CD1E, FLT3, and FCER1A; epithelial cells by EPCAM, KRT18, KRT19, and CDH1; and ciliated cells by epithelial marker expression together with FOXJ1. Dot size represents the percentage of cells expressing each gene, and color intensity indicates the scaled average expression level. (**D**) The expression level of necroptosis-regulatory genes including ZBP1, RIPK1, RIPK3 and MLKL was shown according to scRNA-seq data (n=8 for uninfected control, n=9 for mild cases, and n=7 for fatal cases). Data represented mean and standard error of the mean from the indicated number of subjects. Statistical significance between groups was determined with one way-ANOVA with Tukey’s multiple comparisons tests.

**Fig. S7.**
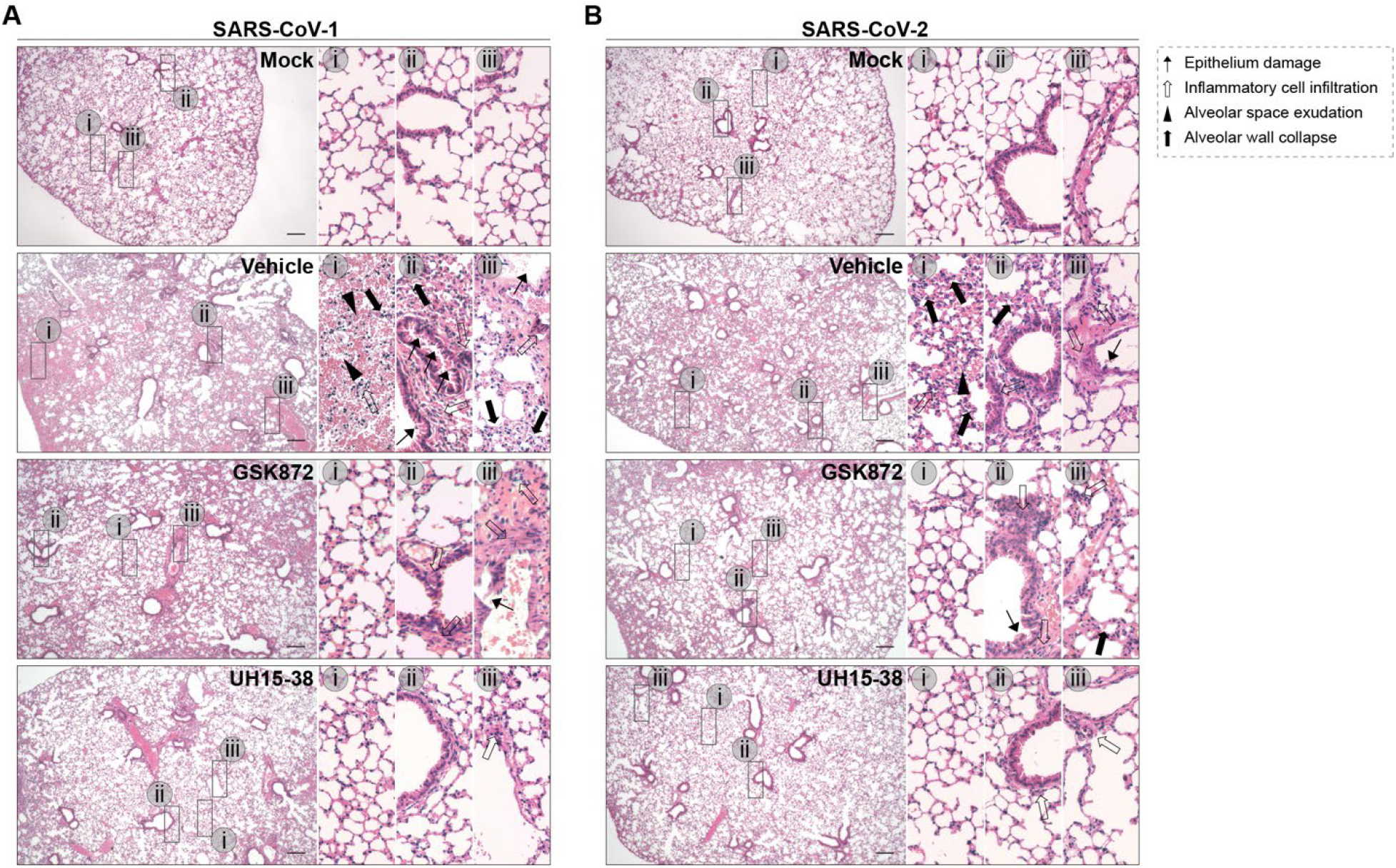
UH15-38 and GSK872 reduce lung histopathological changes induced by SARS-CoV-1 and SARS-CoV-2 infection. (**A-B**) K18-hACE2 mice were inoculated with 1×10^3^ PFU/mouse SARS-CoV-1 or 5×10^3^ PFU/mouse SARS-CoV-2 WT through the i.n. route followed by i.p. administration of 30 mg/kg/day UH15-38, 10 mg/kg/day GSK872, or vehicle for three consecutive days. Animals were sacrificed on 3 dpi. Histopathological changes in the lung were revealed with H&E staining.

**Fig. S8.**
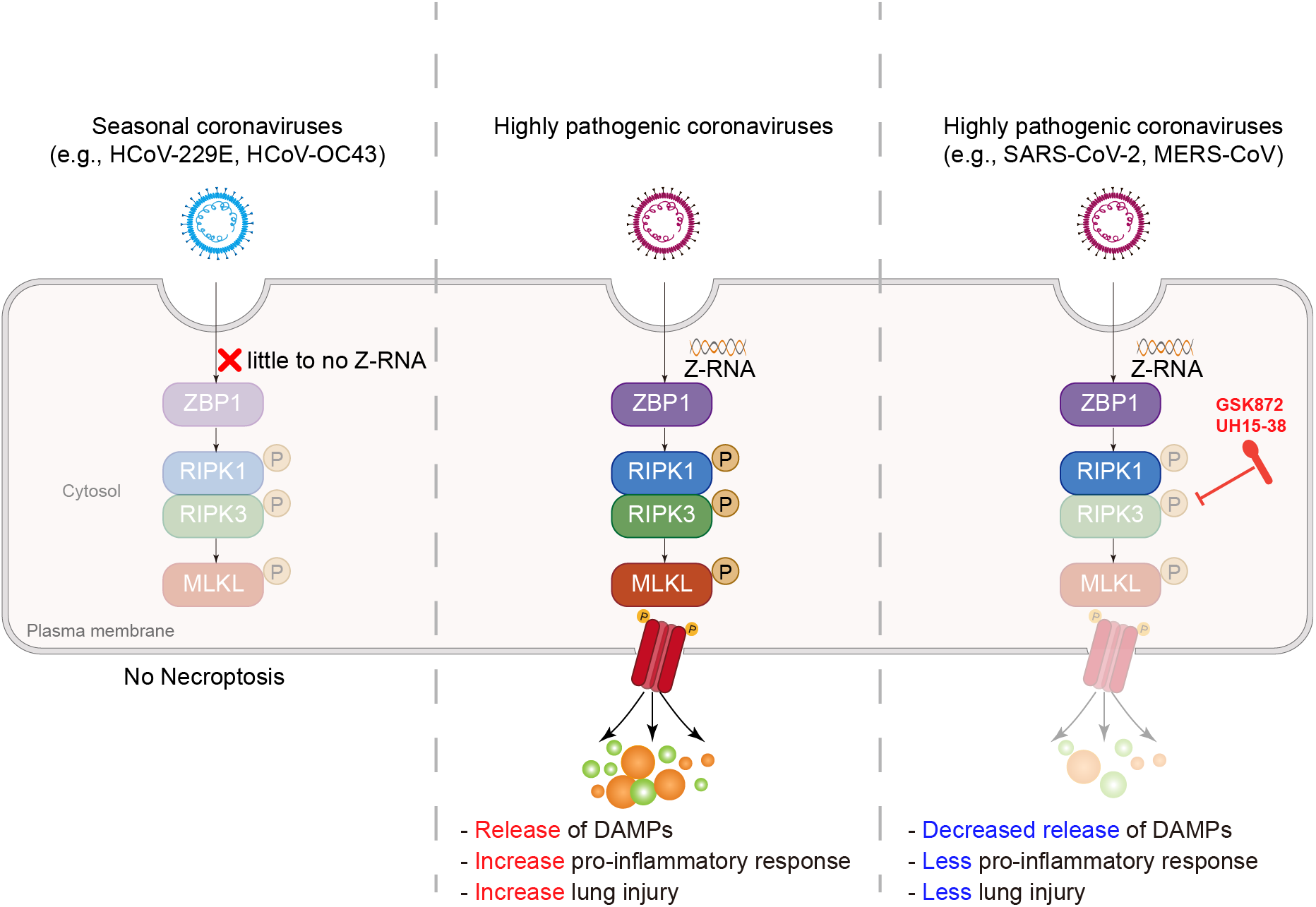
Necroptosis is a pathogenic mechanism of severe coronavirus infection. Schematic of findings of the study.

**Fig. S9.**
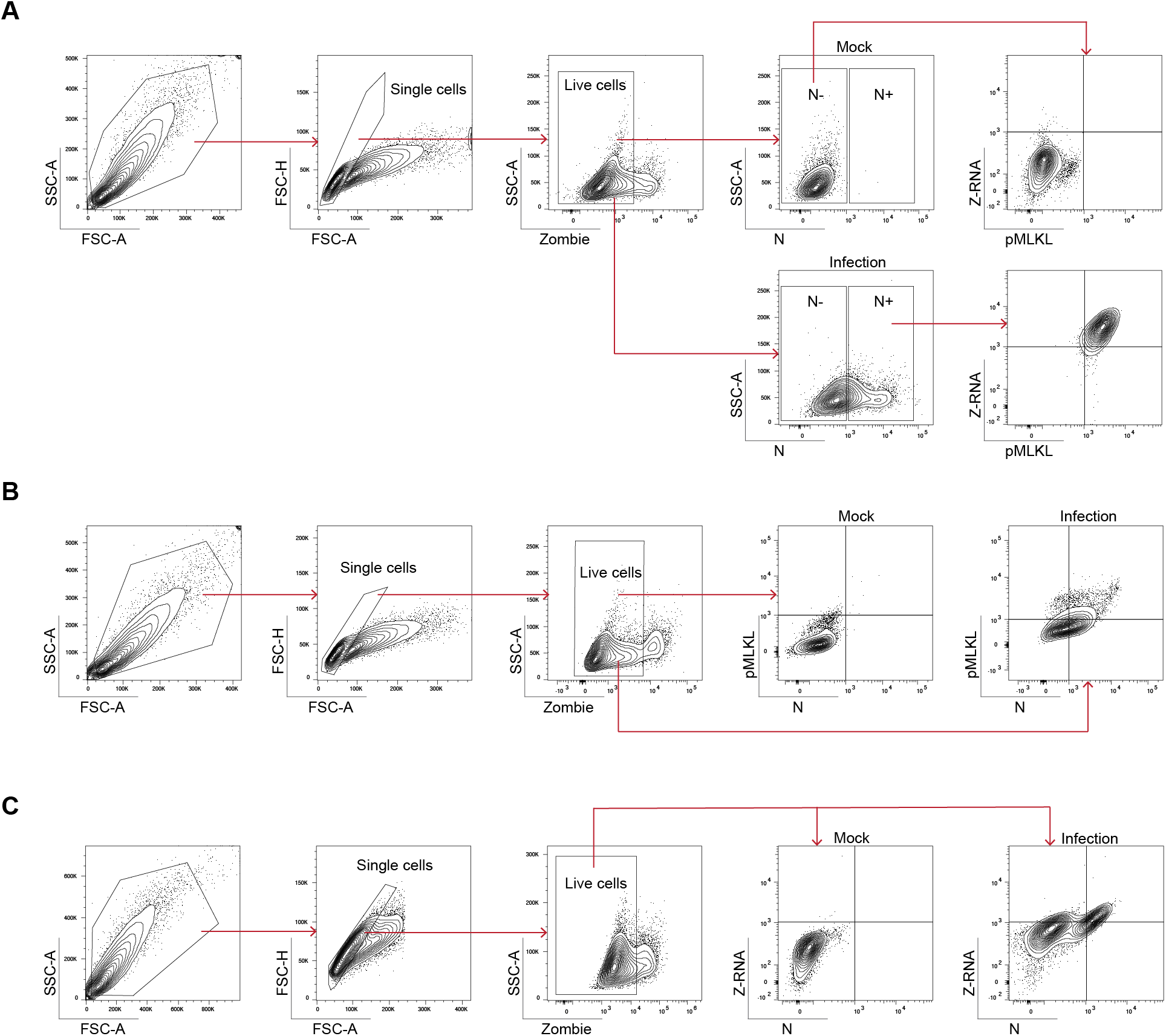
Gating strategies. (**A**) Gating strategy for Fig. S3F. (**B**) Gating strategy for Fig. S4G. (**C**) Gating strategy for Fig. 4F and Fig. S5C.

## References

1. J. F. Chan, S. K. Lau, K. K. To, V. C. Cheng, P. C. Woo, K. Y. Yuen, Middle East respiratory syndrome coronavirus: another zoonotic betacoronavirus causing SARS-like disease. Clin Microbiol Rev 28, 465–522 (2015).

2. J. Cui, F. Li, Z. L. Shi, Origin and evolution of pathogenic coronaviruses. Nat Rev Microbiol 17, 181–192 (2019).

3. P. C. Woo, S. K. Lau, C. S. Lam, C. C. Lau, A. K. Tsang, J. H. Lau, R. Bai, J. L. Teng, C. C. Tsang, M. Wang, B. J. Zheng, K. H. Chan, K. Y. Yuen, Discovery of seven novel Mammalian and avian coronaviruses in the genus deltacoronavirus supports bat coronaviruses as the gene source of alphacoronavirus and betacoronavirus and avian coronaviruses as the gene source of gammacoronavirus and deltacoronavirus. J Virol 86, 3995–4008 (2012).

4. J. A. Lednicky, M. S. Tagliamonte, S. K. White, M. A. Elbadry, M. M. Alam, C. J. Stephenson, T. S. Bonny, J. C. Loeb, T. Telisma, S. Chavannes, D. A. Ostrov, C. Mavian, V. M. Beau De Rochars, M. Salemi, J. G. Morris, Jr., Independent infections of porcine deltacoronavirus among Haitian children. Nature 600, 133–137 (2021).

5. K. Park, M. Shin, A. Natasha, J. Kim, J. Noh, S. G. Kim, B. Kim, J. Park, Y. R. Seo, H. K. Cho, K. S. Byun, J. H. Kim, Y. S. Lee, J. O. Shim, W. K. Kim, J. W. Song, Novel human coronavirus in an infant patient with pneumonia, Republic of Korea. Emerg Microbes Infect 14, 2466705 (2025).

6. A. N. Vlasova, A. Diaz, D. Damtie, L. Xiu, T. H. Toh, J. S. Lee, L. J. Saif, G. C. Gray, Novel Canine Coronavirus Isolated from a Hospitalized Patient With Pneumonia in East Malaysia. Clin Infect Dis 74, 446–454 (2022).

7. A. Zumla, D. S. Hui, M. Peiris, S. Perlman, Respiratory infections due to human common cold coronaviruses, SARS-CoV, MERS-CoV, and SARS-CoV-2: epidemiology, pathogenesis, clinical features, diagnostics, therapeutics, and vaccine landscapes. Lancet Respir Med, (2026).

8. K. O. Alsaad, A. H. Hajeer, M. Al Balwi, M. Al Moaiqel, N. Al Oudah, A. Al Ajlan, S. AlJohani, S. Alsolamy, G. E. Gmati, H. Balkhy, H. H. Al-Jahdali, S. A. Baharoon, Y. M. Arabi, Histopathology of Middle East respiratory syndrome coronovirus (MERS-CoV) infection - clinicopathological and ultrastructural study. Histopathology 72, 516–524 (2018).

9. T. N. Chau, K. C. Lee, H. Yao, T. Y. Tsang, T. C. Chow, Y. C. Yeung, K. W. Choi, Y. K. Tso, T. Lau, S. T. Lai, C. L. Lai, SARS-associated viral hepatitis caused by a novel coronavirus: report of three cases. Hepatology 39, 302–310 (2004).

10. Y. Ding, H. Wang, H. Shen, Z. Li, J. Geng, H. Han, J. Cai, X. Li, W. Kang, D. Weng, Y. Lu, D. Wu, L. He, K. Yao, The clinical pathology of severe acute respiratory syndrome (SARS): a report from China. J Pathol 200, 282–289 (2003).

11. D. L. Ng, F. Al Hosani, M. K. Keating, S. I. Gerber, T. L. Jones, M. G. Metcalfe, S. Tong, Y. Tao, N. N. Alami, L. M. Haynes, M. A. Mutei, L. Abdel-Wareth, T. M. Uyeki, D. L. Swerdlow, M. Barakat, S. R. Zaki, Clinicopathologic, Immunohistochemical, and Ultrastructural Findings of a Fatal Case of Middle East Respiratory Syndrome Coronavirus Infection in the United Arab Emirates, April 2014. Am J Pathol 186, 652–658 (2016).

12. H. Chu, Y. Hou, D. Yang, L. Wen, H. Shuai, C. Yoon, J. Shi, Y. Chai, T. T. Yuen, B. Hu, C. Li, X. Zhao, Y. Wang, X. Huang, K. S. Lee, C. Luo, J. P. Cai, V. K. Poon, C. C. Chan, A. J. Zhang, S. Yuan, K. Y. Sit, D. C. Foo, W. K. Au, K. K. Wong, J. Zhou, K. H. Kok, D. Y. Jin, J. F. Chan, K. Y. Yuen, Coronaviruses exploit a host cysteine-aspartic protease for replication. Nature 609, 785–792 (2022).

13. H. Chu, H. Shuai, Y. Hou, X. Zhang, L. Wen, X. Huang, B. Hu, D. Yang, Y. Wang, C. Yoon, B. H. Wong, C. Li, X. Zhao, V. K. Poon, J. P. Cai, K. K. Wong, M. L. Yeung, J. Zhou, R. K. Au-Yeung, S. Yuan, D. Y. Jin, K. H. Kok, S. Perlman, J. F. Chan, K. Y. Yuen, Targeting highly pathogenic coronavirus-induced apoptosis reduces viral pathogenesis and disease severity. Sci Adv 7, (2021).

14. A. C. Sims, H. D. Mitchell, L. E. Gralinski, J. E. Kyle, K. E. Burnum-Johnson, M. Lam, M. L. Fulcher, A. West, R. D. Smith, S. H. Randell, T. O. Metz, T. P. Sheahan, K. M. Waters, R. S. Baric, Unfolded Protein Response Inhibition Reduces Middle East Respiratory Syndrome Coronavirus-Induced Acute Lung Injury. mBio 12, e0157221 (2021).

15. A. Degterev, Z. Huang, M. Boyce, Y. Li, P. Jagtap, N. Mizushima, G. D. Cuny, T. J. Mitchison, M. A. Moskowitz, J. Yuan, Chemical inhibitor of nonapoptotic cell death with therapeutic potential for ischemic brain injury. Nat Chem Biol 1, 112–119 (2005).

16. D. Vercammen, R. Beyaert, G. Denecker, V. Goossens, G. Van Loo, W. Declercq, J. Grooten, W. Fiers, P. Vandenabeele, Inhibition of caspases increases the sensitivity of L929 cells to necrosis mediated by tumor necrosis factor. J Exp Med 187, 1477–1485 (1998).

17. Y. S. Cho, S. Challa, D. Moquin, R. Genga, T. D. Ray, M. Guildford, F. K. Chan, Phosphorylation-driven assembly of the RIP1-RIP3 complex regulates programmed necrosis and virus-induced inflammation. Cell 137, 1112–1123 (2009).

18. F. K. Chan, J. Shisler, J. G. Bixby, M. Felices, L. Zheng, M. Appel, J. Orenstein, B. Moss, M. J. Lenardo, A role for tumor necrosis factor receptor-2 and receptor-interacting protein in programmed necrosis and antiviral responses. J Biol Chem 278, 51613–51621 (2003).

19. N. Holler, R. Zaru, O. Micheau, M. Thome, A. Attinger, S. Valitutti, J. L. Bodmer, P. Schneider, B. Seed, J. Tschopp, Fas triggers an alternative, caspase-8-independent cell death pathway using the kinase RIP as effector molecule. Nat Immunol 1, 489–495 (2000).

20. H. Guo, S. Omoto, P. A. Harris, J. N. Finger, J. Bertin, P. J. Gough, W. J. Kaiser, E. S. Mocarski, Herpes simplex virus suppresses necroptosis in human cells. Cell Host Microbe 17, 243–251 (2015).

21. Z. Huang, S. Q. Wu, Y. Liang, X. Zhou, W. Chen, L. Li, J. Wu, Q. Zhuang, C. Chen, J. Li, C. Q. Zhong, W. Xia, R. Zhou, C. Zheng, J. Han, RIP1/RIP3 binding to HSV-1 ICP6 initiates necroptosis to restrict virus propagation in mice. Cell Host Microbe 17, 229–242 (2015).

22. J. W. Upton, W. J. Kaiser, E. S. Mocarski, Virus inhibition of RIP3-dependent necrosis. Cell Host Microbe 7, 302–313 (2010).

23. J. W. Upton, W. J. Kaiser, E. S. Mocarski, DAI/ZBP1/DLM-1 complexes with RIP3 to mediate virus-induced programmed necrosis that is targeted by murine cytomegalovirus vIRA. Cell Host Microbe 11, 290–297 (2012).

24. H. Koehler, S. Cotsmire, T. Zhang, S. Balachandran, J. W. Upton, J. Langland, D. Kalman, B. L. Jacobs, E. S. Mocarski, Vaccinia virus E3 prevents sensing of Z-RNA to block ZBP1-dependent necroptosis. Cell Host Microbe 29, 1266–1276 e1265 (2021).

25. S. Nogusa, R. J. Thapa, C. P. Dillon, S. Liedmann, T. H. Oguin, 3rd, J. P. Ingram, D. A. Rodriguez, R. Kosoff, S. Sharma, O. Sturm, K. Verbist, P. J. Gough, J. Bertin, B. M. Hartmann, S. C. Sealfon, W. J. Kaiser, E. S. Mocarski, C. B. Lopez, P. G. Thomas, A. Oberst, D. R. Green, S. Balachandran, RIPK3 Activates Parallel Pathways of MLKL-Driven Necroptosis and FADD-Mediated Apoptosis to Protect against Influenza A Virus. Cell Host Microbe 20, 13–24 (2016).

26. R. J. Thapa, J. P. Ingram, K. B. Ragan, S. Nogusa, D. F. Boyd, A. A. Benitez, H. Sridharan, R. Kosoff, M. Shubina, V. J. Landsteiner, M. Andrake, P. Vogel, L. J. Sigal, B. R. tenOever, P. G. Thomas, J. W. Upton, S. Balachandran, DAI Senses Influenza A Virus Genomic RNA and Activates RIPK3-Dependent Cell Death. Cell Host Microbe 20, 674–681 (2016).

27. T. Zhang, C. Yin, D. F. Boyd, G. Quarato, J. P. Ingram, M. Shubina, K. B. Ragan, T. Ishizuka, J. C. Crawford, B. Tummers, D. A. Rodriguez, J. Xue, S. Peri, W. J. Kaiser, C. B. Lopez, Y. Xu, J. W. Upton, P. G. Thomas, D. R. Green, S. Balachandran, Influenza Virus Z-RNAs Induce ZBP1-Mediated Necroptosis. Cell 180, 1115–1129 e1113 (2020).

28. T. Kuriakose, S. M. Man, R. K. Malireddi, R. Karki, S. Kesavardhana, D. E. Place, G. Neale, P. Vogel, T. D. Kanneganti, ZBP1/DAI is an innate sensor of influenza virus triggering the NLRP3 inflammasome and programmed cell death pathways. Sci Immunol 1, (2016).

29. S. Li, Y. Zhang, Z. Guan, H. Li, M. Ye, X. Chen, J. Shen, Y. Zhou, Z. L. Shi, P. Zhou, K. Peng, SARS-CoV-2 triggers inflammatory responses and cell death through caspase-8 activation. Signal Transduct Target Ther 5, 235 (2020).

30. K. Liang, K. C. Barnett, M. Hsu, W. C. Chou, S. S. Bais, K. Riebe, Y. Xie, T. T. Nguyen, T. H. Oguin, 3rd, K. M. Vannella, S. M. Hewitt, D. S. Chertow, M. Blasi, G. D. Sempowski, A. Karlsson, B. H. Koller, D. J. Lenschow, S. H. Randell, J. P. Ting, Initiator cell death event induced by SARS-CoV-2 in the human airway epithelium. Sci Immunol 9, eadn0178 (2024).

31. A. Elsharkawy, H. R. Jahantigh, A. Guglani, S. Stone, K. Arora, M. Kumar, Virus-specific host responses and gene signatures following infection with major SARS-CoV-2 variants of concern: role of ZBP1 in viral clearance and lung inflammation. Front Immunol 16, 1557535 (2025).

32. M. Evdokimova, S. Feng, A. Caobi, F. R. Moreira, D. Jones, K. D. Alysandratos, E. S. Tully, D. N. Kotton, D. F. Boyd, B. S. Banach, R. N. Kirchdoerfer, M. Saeed, S. C. Baker, Coronavirus endoribonuclease antagonizes ZBP1-mediated necroptosis and delays multiple cell death pathways. Proc Natl Acad Sci U S A 122, e2419620122 (2025).

33. S. Li, Y. Zhang, Z. Guan, M. Ye, H. Li, M. You, Z. Zhou, C. Zhang, F. Zhang, B. Lu, P. Zhou, K. Peng, SARS-CoV-2 Z-RNA activates the ZBP1-RIPK3 pathway to promote virus-induced inflammatory responses. Cell Res 33, 201–214 (2023).

34. H. Wang, M. Liang, J. Zhang, H. Tong, F. Zhang, Y. Liu, P. Wang, M. Chang, F. Han, S. Liu, Y. Lin, W. Song, R. Karki, P. Wang, H. Chen, Y. Liu, M. Zheng, The NSP5, ORF6 and NSP13 of SARS-CoV-2 Cooperate to Modulate Inflammatory Cell Death Activation. Adv Sci (Weinh) 12, e03977 (2025).

35. M. B. S, J. P. Cooney, R. Bhandari, L. Mackiewicz, M. Dayton, D. Sheerin, S. R. Georgy, J. M. Murphy, K. C. Davidson, C. C. Allison, M. Pellegrini, M. Doerflinger, Necroptosis does not drive disease pathogenesis in a mouse infective model of SARS-CoV-2 in vivo. Cell Death Dis 15, 100 (2024).

36. H. Nailwal, F. K. Chan, Necroptosis in anti-viral inflammation. Cell Death Differ 26, 4–13 (2019).

37. F. K. Chan, Fueling the flames: Mammalian programmed necrosis in inflammatory diseases. Cold Spring Harb Perspect Biol 4, (2012).

38. K. Moriwaki, J. Bertin, P. J. Gough, G. M. Orlowski, F. K. Chan, Differential roles of RIPK1 and RIPK3 in TNF-induced necroptosis and chemotherapeutic agent-induced cell death. Cell Death Dis 6, e1636 (2015).

39. G. B. Koo, M. J. Morgan, D. G. Lee, W. J. Kim, J. H. Yoon, J. S. Koo, S. I. Kim, S. J. Kim, M. K. Son, S. S. Hong, J. M. Levy, D. A. Pollyea, C. T. Jordan, P. Yan, D. Frankhouser, D. Nicolet, K. Maharry, G. Marcucci, K. S. Choi, H. Cho, A. Thorburn, Y. S. Kim, Methylation-dependent loss of RIP3 expression in cancer represses programmed necrosis in response to chemotherapeutics. Cell Res 25, 707–725 (2015).

40. P. G. Thomas, M. Shubina, S. Balachandran, ZBP1/DAI-Dependent Cell Death Pathways in Influenza A Virus Immunity and Pathogenesis. Curr Top Microbiol Immunol 442, 41–63 (2023).

41. H. Hofmann, K. Pyrc, L. van der Hoek, M. Geier, B. Berkhout, S. Pohlmann, Human coronavirus NL63 employs the severe acute respiratory syndrome coronavirus receptor for cellular entry. Proc Natl Acad Sci U S A 102, 7988–7993 (2005).

42. J. Maelfait, L. Liverpool, A. Bridgeman, K. B. Ragan, J. W. Upton, J. Rehwinkel, Sensing of viral and endogenous RNA by ZBP1/DAI induces necroptosis. EMBO J 36, 2529–2543 (2017).

43. P. S. Ho, M. J. Ellison, G. J. Quigley, A. Rich, A computer aided thermodynamic approach for predicting the formation of Z-DNA in naturally occurring sequences. EMBO J 5, 2737–2744 (1986).

44. P. S. Ho, Thermogenomics: thermodynamic-based approaches to genomic analyses of DNA structure. Methods 47, 159–167 (2009).

45. H. Shuai, J. F. Chan, B. Hu, Y. Chai, C. Yoon, H. Liu, Y. Liu, J. Shi, T. Zhu, J. C. Hu, Y. F. Hu, Y. Hou, X. Huang, T. T. Yuen, Y. Wang, J. Zhang, Y. Xia, L. L. Chen, J. P. Cai, A. J. Zhang, S. Yuan, J. Zhou, B. Z. Zhang, J. D. Huang, K. Y. Yuen, K. K. To, H. Chu, The viral fitness and intrinsic pathogenicity of dominant SARS-CoV-2 Omicron sublineages BA.1, BA.2, and BA.5. EBioMedicine 95, 104753 (2023).

46. H. Shuai, J. F. Chan, B. Hu, Y. Chai, T. T. Yuen, F. Yin, X. Huang, C. Yoon, J. C. Hu, H. Liu, J. Shi, Y. Liu, T. Zhu, J. Zhang, Y. Hou, Y. Wang, L. Lu, J. P. Cai, A. J. Zhang, J. Zhou, S. Yuan, M. A. Brindley, B. Z. Zhang, J. D. Huang, K. K. To, K. Y. Yuen, H. Chu, Attenuated replication and pathogenicity of SARS-CoV-2 B.1.1.529 Omicron. Nature 603, 693–699 (2022).

47. K. Li, C. L. Wohlford-Lenane, R. Channappanavar, J. E. Park, J. T. Earnest, T. B. Bair, A. M. Bates, K. A. Brogden, H. A. Flaherty, T. Gallagher, D. K. Meyerholz, S. Perlman, P. B. McCray, Jr., Mouse-adapted MERS coronavirus causes lethal lung disease in human DPP4 knockin mice. Proc Natl Acad Sci U S A 114, E3119–E3128 (2017).

48. P. Mandal, S. B. Berger, S. Pillay, K. Moriwaki, C. Huang, H. Guo, J. D. Lich, J. Finger, V. Kasparcova, B. Votta, M. Ouellette, B. W. King, D. Wisnoski, A. S. Lakdawala, M. P. DeMartino, L. N. Casillas, P. A. Haile, C. A. Sehon, R. W. Marquis, J. Upton, L. P. Daley-Bauer, L. Roback, N. Ramia, C. M. Dovey, J. E. Carette, F. K. Chan, J. Bertin, P. J. Gough, E. S. Mocarski, W. J. Kaiser, RIP3 induces apoptosis independent of pronecrotic kinase activity. Mol Cell 56, 481–495 (2014).

49. A. Gautam, D. F. Boyd, S. Nikhar, T. Zhang, I. Siokas, L. A. Van de Velde, J. Gaevert, V. Meliopoulos, B. Thapa, D. A. Rodriguez, K. Q. Cai, C. Yin, D. Schnepf, J. Beer, C. DeAntoneo, R. M. Williams, M. Shubina, B. Livingston, D. Zhang, M. D. Andrake, S. Lee, R. Boda, A. L. Duddupudi, J. C. Crawford, P. Vogel, C. Loch, M. Schwemmle, L. C. Fritz, S. Schultz-Cherry, D. R. Green, G. D. Cuny, P. G. Thomas, A. Degterev, S. Balachandran, Necroptosis blockade prevents lung injury in severe influenza. Nature 628, 835–843 (2024).

50. Z. Liu, H. Nailwal, J. Rector, M. M. Rahman, R. Sam, G. McFadden, F. K. Chan, A class of viral inducer of degradation of the necroptosis adaptor RIPK3 regulates virus-induced inflammation. Immunity 54, 247–258 e247 (2021).

51. A. K. Reuschl, L. G. Thorne, M. V. X. Whelan, R. Ragazzini, W. Furnon, V. M. Cowton, G. De Lorenzo, D. Mesner, J. L. E. Turner, G. Dowgier, N. Bogoda, P. Bonfanti, M. Palmarini, A. H. Patel, C. Jolly, G. J. Towers, Evolution of enhanced innate immune suppression by SARS-CoV-2 Omicron subvariants. Nat Microbiol 9, 451–463 (2024).

52. A. Wickenhagen, M. Flagg, J. R. Port, C. K. Yinda, K. Goldin, S. Gallogly, J. E. Schulz, T. Lutterman, B. N. Williamson, F. Kaiser, R. K. Mukesh, S. van Tol, B. Smith, N. van Doremalen, C. A. Russell, E. de Wit, V. J. Munster, Evolution of Omicron lineage towards increased fitness in the upper respiratory tract in the absence of severe lung pathology. Nat Commun 16, 594 (2025).

53. C. J. Otter, D. M. Renner, A. Fausto, L. H. Tan, N. A. Cohen, S. R. Weiss, Interferon signaling in the nasal epithelium distinguishes among lethal and common cold coronaviruses and mediates viral clearance. Proc Natl Acad Sci U S A 121, e2402540121 (2024).

54. C. J. Otter, A. Fausto, L. H. Tan, A. S. Khosla, N. A. Cohen, S. R. Weiss, Infection of primary nasal epithelial cells differentiates among lethal and seasonal human coronaviruses. Proc Natl Acad Sci U S A 120, e2218083120 (2023).

55. T. I. Breugem, S. Riesebosch, J. Zhang, A. Z. Mykytyn, L. Krabbendam, N. Groen, S. Baptista Varela, D. Schipper, P. B. van den Doel, R. van Acker, R. Stadhouders, M. M. Lamers, B. L. Haagmans, Variable DPP4 expression in multiciliated cells of the human nasal epithelium as a determinant for MERS-CoV tropism. Proc Natl Acad Sci U S A 122, e2410630122 (2025).

56. B. Rockx, T. Kuiken, S. Herfst, T. Bestebroer, M. M. Lamers, B. B. Oude Munnink, D. de Meulder, G. van Amerongen, J. van den Brand, N. M. A. Okba, D. Schipper, P. van Run, L. Leijten, R. Sikkema, E. Verschoor, B. Verstrepen, W. Bogers, J. Langermans, C. Drosten, M. Fentener van Vlissingen, R. Fouchier, R. de Swart, M. Koopmans, B. L. Haagmans, Comparative pathogenesis of COVID-19, MERS, and SARS in a nonhuman primate model. Science 368, 1012–1015 (2020).

57. J. Prescott, D. Falzarano, E. de Wit, K. Hardcastle, F. Feldmann, E. Haddock, D. Scott, H. Feldmann, V. J. Munster, Pathogenicity and Viral Shedding of MERS-CoV in Immunocompromised Rhesus Macaques. Front Immunol 9, 205 (2018).

58. Y. Dondelinger, P. Hulpiau, Y. Saeys, M. J. M. Bertrand, P. Vandenabeele, An evolutionary perspective on the necroptotic pathway. Trends Cell Biol 26, 721–732 (2016).

59. A. Agueda-Pinto, L. Q. Alves, F. Neves, G. McFadden, B. L. Jacobs, L. F. C. Castro, M. M. Rahman, P. J. Esteves, Convergent Loss of the Necroptosis Pathway in Disparate Mammalian Lineages Shapes Viruses Countermeasures. Front Immunol 12, 747737 (2021).

60. M. Ahn, D. E. Anderson, Q. Zhang, C. W. Tan, B. L. Lim, K. Luko, M. Wen, W. N. Chia, S. Mani, L. C. Wang, J. H. J. Ng, R. M. Sobota, C. A. Dutertre, F. Ginhoux, Z. L. Shi, A. T. Irving, L. F. Wang, Dampened NLRP3-mediated inflammation in bats and implications for a special viral reservoir host. Nat Microbiol 4, 789–799 (2019).

61. S. Das, D. Jain, P. Chaudhary, R. M. Quintela-Tizon, A. Banerjee, S. Kesavardhana, Bat adaptations in inflammation and cell death regulation contribute to viral tolerance. mBio 16, e0320423 (2025).

62. M. Ahn, V. C. Chen, P. Rozario, W. L. Ng, P. S. Kong, W. R. Sia, A. E. Z. Kang, Q. Su, L. H. Nguyen, F. Zhu, W. O. Y. Chan, C. W. Tan, W. S. Cheong, Y. Y. Hey, R. Foo, F. Guo, Y. T. Lim, X. Li, W. N. Chia, R. M. Sobota, N. Y. Fu, A. T. Irving, L. F. Wang, Bat ASC2 suppresses inflammasomes and ameliorates inflammatory diseases. Cell 186, 2144–2159 e2122 (2023).

63. J. Xie, Y. Li, X. Shen, G. Goh, Y. Zhu, J. Cui, L. F. Wang, Z. L. Shi, P. Zhou, Dampened STING-Dependent Interferon Activation in Bats. Cell Host Microbe 23, 297–301 e294 (2018).

64. C. Yin, A. Fedorov, H. Guo, J. C. Crawford, C. Rousseau, X. Zhong, R. M. Williams, A. Gautam, H. S. Koehler, A. W. Whisnant, T. Hennig, A. Rozina, Y. Zhong, S. Lv, V. Bergant, S. Wang, P. Droge, S. Miller, M. Poptsova, J. Rehwinkel, A. Pichlmair, E. S. Mocarski, P. G. Thomas, L. Dolken, T. Zhang, A. Herbert, S. Balachandran, Host cell Z-RNAs activate ZBP1 during virus infections. Nature 648, 707–716 (2025).

65. P. C. Woo, M. Wang, S. K. Lau, H. Xu, R. W. Poon, R. Guo, B. H. Wong, K. Gao, H. W. Tsoi, Y. Huang, K. S. Li, C. S. Lam, K. H. Chan, B. J. Zheng, K. Y. Yuen, Comparative analysis of twelve genomes of three novel group 2c and group 2d coronaviruses reveals unique group and subgroup features. J Virol 81, 1574–1585 (2007).

66. H. Chu, C. M. Chan, X. Zhang, Y. Wang, S. Yuan, J. Zhou, R. K. Au-Yeung, K. H. Sze, D. Yang, H. Shuai, Y. Hou, C. Li, X. Zhao, V. K. Poon, S. P. Leung, M. L. Yeung, J. Yan, G. Lu, D. Y. Jin, G. F. Gao, J. F. Chan, K. Y. Yuen, Middle East respiratory syndrome coronavirus and bat coronavirus HKU9 both can utilize GRP78 for attachment onto host cells. J Biol Chem 293, 11709–11726 (2018).

67. J. F. Chan, K. H. Chan, G. K. Choi, K. K. To, H. Tse, J. P. Cai, M. L. Yeung, V. C. Cheng, H. Chen, X. Y. Che, S. K. Lau, P. C. Woo, K. Y. Yuen, Differential cell line susceptibility to the emerging novel human betacoronavirus 2c EMC/2012: implications for disease pathogenesis and clinical manifestation. J Infect Dis 207, 1743–1752 (2013).

68. A. M. Zaki, S. van Boheemen, T. M. Bestebroer, A. D. Osterhaus, R. A. Fouchier, Isolation of a novel coronavirus from a man with pneumonia in Saudi Arabia. N Engl J Med 367, 1814–1820 (2012).

69. H. Chu, J. F. Chan, T. T. Yuen, H. Shuai, S. Yuan, Y. Wang, B. Hu, C. C. Yip, J. O. Tsang, X. Huang, Y. Chai, D. Yang, Y. Hou, K. K. Chik, X. Zhang, A. Y. Fung, H. W. Tsoi, J. P. Cai, W. M. Chan, J. D. Ip, A. W. Chu, J. Zhou, D. C. Lung, K. H. Kok, K. K. To, O. T. Tsang, K. H. Chan, K. Y. Yuen, Comparative tropism, replication kinetics, and cell damage profiling of SARS-CoV-2 and SARS-CoV with implications for clinical manifestations, transmissibility, and laboratory studies of COVID-19: an observational study. Lancet Microbe 1, e14–e23 (2020).

70. H. Shuai, J. Qiao, C. Yoon, G. Zhang, Y. Hou, X. Xia, L. Wang, X. Deng, Y. Wang, Q. Li, L. Du, Y. Liu, M. Zhou, H. T. Wong, J. Hu, H. Liu, B. Hu, D. Wang, J. Su, Y. Pan, Y. Ye, Y. Chen, Z. Fang, Z. Xia, Y. Chai, J. Shi, Y. Wang, T. Zhu, H. Zhang, S. Yuan, J. Zhou, J. F. Chan, K. Y. Yuen, C. Xu, J. Lei, S. Yang, H. Chu, An orally available M(pro)/TMPRSS2 bispecific inhibitor with potent anti-coronavirus efficacy in vivo. Nat Commun 16, 6541 (2025).

71. J. F. Chan, Y. J. Oh, S. Yuan, H. Chu, M. L. Yeung, D. Canena, C. C. Chan, V. K. Poon, C. C. Chan, A. J. Zhang, J. P. Cai, Z. W. Ye, L. Wen, T. T. Yuen, K. K. Chik, H. Shuai, Y. Wang, Y. Hou, C. Luo, W. M. Chan, Z. Qin, K. Y. Sit, W. K. Au, M. Legendre, R. Zhu, L. Hain, H. Seferovic, R. Tampe, K. K. To, K. H. Chan, D. G. Thomas, M. Klausberger, C. Xu, J. J. Moon, J. Stadlmann, J. M. Penninger, C. Oostenbrink, P. Hinterdorfer, K. Y. Yuen, D. M. Markovitz, A molecularly engineered, broad-spectrum anti-coronavirus lectin inhibits SARS-CoV-2 and MERS-CoV infection in vivo. Cell Rep Med 3, 100774 (2022).

72. H. Chu, J. F. Chan, Y. Wang, T. T. Yuen, Y. Chai, Y. Hou, H. Shuai, D. Yang, B. Hu, X. Huang, X. Zhang, J. P. Cai, J. Zhou, S. Yuan, K. H. Kok, K. K. To, I. H. Chan, A. J. Zhang, K. Y. Sit, W. K. Au, K. Y. Yuen, Comparative Replication and Immune Activation Profiles of SARS-CoV-2 and SARS-CoV in Human Lungs: An Ex Vivo Study With Implications for the Pathogenesis of COVID-19. Clin Infect Dis 71, 1400–1409 (2020).

73. T. Sato, D. E. Stange, M. Ferrante, R. G. Vries, J. H. Van Es, S. Van den Brink, W. J. Van Houdt, A. Pronk, J. Van Gorp, P. D. Siersema, H. Clevers, Long-term expansion of epithelial organoids from human colon, adenoma, adenocarcinoma, and Barrett’s epithelium. Gastroenterology 141, 1762–1772 (2011).

74. J. Zhou, C. Li, G. Zhao, H. Chu, D. Wang, H. H. Yan, V. K. Poon, L. Wen, B. H. Wong, X. Zhao, M. C. Chiu, D. Yang, Y. Wang, R. K. H. Au-Yeung, I. H. Chan, S. Sun, J. F. Chan, K. K. To, Z. A. Memish, V. M. Corman, C. Drosten, I. F. Hung, Y. Zhou, S. Y. Leung, K. Y. Yuen, Human intestinal tract serves as an alternative infection route for Middle East respiratory syndrome coronavirus. Sci Adv 3, eaao4966 (2017).

75. X. Zhao, C. Li, X. Liu, M. C. Chiu, D. Wang, Y. Wei, H. Chu, J. P. Cai, I. Hau-Yee Chan, K. Kak-Yuen Wong, J. Fuk-Woo Chan, K. Kai-Wang To, K. Y. Yuen, J. Zhou, Human Intestinal Organoids Recapitulate Enteric Infections of Enterovirus and Coronavirus. Stem Cell Reports 16, 493–504 (2021).

76. Y. Liu, X. Zhao, J. Shi, Y. Wang, H. Liu, Y. F. Hu, B. Hu, H. Shuai, T. T. Yuen, Y. Chai, F. Liu, H. R. Gong, J. Li, X. Wang, S. Jiang, X. Zhang, Y. Zhang, X. Li, L. Wang, M. Hartnoll, T. Zhu, Y. Hou, X. Huang, C. Yoon, Y. Wang, Y. He, M. Zhou, L. Du, X. Zhang, W. M. Chan, L. L. Chen, J. P. Cai, S. Yuan, J. Zhou, J. D. Huang, K. Y. Yuen, K. K. To, J. F. Chan, B. Z. Zhang, L. Sun, P. Wang, H. Chu, Lineage-specific pathogenicity, immune evasion, and virological features of SARS-CoV-2 BA.2.86/JN.1 and EG.5.1/HK.3. Nat Commun 15, 8728 (2024).

77. J. Shi, X. Zhao, X. Jin, J. Li, Y. Liu, H. Liu, Y. F. Hu, Z. Chen, Y. Xiao, L. Wang, Y. Wang, Y. He, Y. Chai, B. Hu, H. Shuai, Y. Wang, X. Li, S. Jiang, Y. Zhang, X. Zhang, W. M. Chan, L. L. Chen, J. P. Cai, B. Sui, H. Zhang, D. Yang, L. Zhu, S. Yuan, J. Zhou, J. D. Huang, K. Y. Yuen, K. K. To, J. F. Chan, B. Z. Zhang, Q. Wang, M. He, L. Sun, P. Wang, H. Chu, Pathogenicity, virological features, and immune evasion of SARS-CoV-2 JN.1-derived variants including JN.1.7, KP.2, KP.3, and KP.3.1.1. Nat Commun 16, 11002 (2025).

67. J. F. Chan et al., Differential cell line susceptibility to the emerging novel human betacoronavirus 2c EMC/2012: implications for disease pathogenesis and clinical manifestation. J Infect Dis 207, 1743–1752 (2013).

69. H. Chu et al., Comparative tropism, replication kinetics, and cell damage profiling of SARS-CoV-2 and SARS-CoV with implications for clinical manifestations, transmissibility, and laboratory studies of COVID-19: an observational study. Lancet Microbe 1, e14–e23 (2020).

70. H. Shuai et al., An orally available M(pro)/TMPRSS2 bispecific inhibitor with potent anti-coronavirus efficacy in vivo. Nat Commun 16, 6541 (2025).

71. J. F. Chan et al., A molecularly engineered, broad-spectrum anti-coronavirus lectin inhibits SARS-CoV-2 and MERS-CoV infection in vivo. Cell Rep Med 3, 100774 (2022).

72. H. Chu et al., Comparative Replication and Immune Activation Profiles of SARS-CoV-2 and SARS-CoV in Human Lungs: An Ex Vivo Study With Implications for the Pathogenesis of COVID-19. Clin Infect Dis 71, 1400–1409 (2020).

73. T. Sato et al., Long-term expansion of epithelial organoids from human colon, adenoma, adenocarcinoma, and Barrett’s epithelium. Gastroenterology 141, 1762–1772 (2011).

74. J. Zhou et al., Human intestinal tract serves as an alternative infection route for Middle East respiratory syndrome coronavirus. Sci Adv 3, eaao4966 (2017).

75. X. Zhao et al., Human Intestinal Organoids Recapitulate Enteric Infections of Enterovirus and Coronavirus. Stem Cell Reports 16, 493–504 (2021).

76. Y. Liu et al., Lineage-specific pathogenicity, immune evasion, and virological features of SARS-CoV-2 BA.2.86/JN.1 and EG.5.1/HK.3. Nat Commun 15, 8728 (2024).

77. J. Shi et al., Pathogenicity, virological features, and immune evasion of SARS-CoV-2 JN.1-derived variants including JN.1.7, KP.2, KP.3, and KP.3.1.1. Nat Commun 16, 11002 (2025).

